# Evaluating Single-Cell Perturbation Response Models Is Far from Straightforward

**DOI:** 10.64898/2026.02.14.705879

**Authors:** Mahshid Heidari, Mina Karimpour, Sumana Srivatsa, Hesam Montazeri

## Abstract

Predicting cellular responses to genetic and chemical perturbations remains a central challenge in single-cell biology and a key step toward building *in silico* virtual cells. The rapid growth of perturbation datasets and advances in deep-learning models have raised expectations for accurate and generalizable prediction. We show that these expectations are overly optimistic, largely due to the failure modes of existing evaluation metrics.

In this study, using cross-splitting, controlled noise experiments, and synthetic data, we systematically evaluate both prediction models and evaluation metrics. We demonstrate that widely used metrics, including correlation-based measures and common distributional distances, are strongly influenced by scale, sparsity, and dimensionality, often misrepresenting model performance. In particular, the Wasserstein distance fails in high-dimensional gene expression spaces under variance scaling, while the Energy distance can overlook disruptions in gene-gene dependencies. Our analyses further reveal that complex deep learning models often underperform simple baselines and remain far from empirical performance bounds across multiple chemical perturbation datasets. Together, our framework exposes critical pitfalls, establishes robust evaluation guidelines, and provides a foundation for trustworthy benchmarking toward reliable virtual-cell models.

## Introduction

Single-cell transcriptomics technologies have transformed molecular biology by providing unprecedented insights into cellular heterogeneity, dynamic and transcriptomic complexity^1,2^. One notable application is in perturbation experiments, which explore treatment effects to guide personalized therapies, however, such experiments are resource-intensive, expensive and not universally applicable to all cells or tissues^3^.

Therefore, a wide range of computational approaches has emerged to predict unseen genetic or chemical perturbation responses using available single-cell RNA sequencing (scRNA-seq) perturbation datasets, spanning diverse architectural paradigms. These include variational autoencoder-based generative models^4^, graph-based methods leveraging gene-gene interaction networks^5^, optimal transport-based frameworks for modeling cell-state transitions^6,7^, and more recent foundation models that adopt large-scale pretraining and large language model-inspired architectures^8,9^. Together, these approaches reflect the growing methodological diversity aimed at capturing complex regulatory programs and predicting cellular responses under perturbation.

However, accurately predicting perturbation responses at single-cell resolution remains challenging, and several studies have claimed that complex model architectures often provide little or no improvement over simple baselines. In particular, the no-perturbation model, which assumes that perturbations have no impact on gene expression, as well as simple multilayer perceptron (MLP) models, have frequently been shown to achieve performance comparable to that of more complex models^10,11^. In addition, recent evaluations of large single-cell foundation models including scGPT, scFoundation, scBERT^12^ and Geneformer^13^ have reported that such complex models often fail to outperform simpler linear baseline models in perturbation prediction tasks^11,14–17^.

At the same time, emerging work has highlighted fundamental limitations of the benchmarking metrics themselves. Viñas Torné *et al*. showed that commonly used average-based evaluation metrics can substantially overestimate model performance in genetic perturbation datasets due to systematic variation and confounding effects^18^. Miller *et al*., using positive and negative controls and defining a measure of *calibration*, argued that deep learning-based genetic perturbation models can outperform uninformative baselines when evaluated on well-calibrated average-based metrics^19,20^, emphasizing that conclusions about model effectiveness are highly sensitive to the choice of evaluation criteria.

Together, these observations underscore a critical challenge in the field: the need for evaluation criteria and benchmarking frameworks that not only accurately measure predictive power relative to null models, but also assess how well models capture the intrinsic heterogeneity of single-cell data. Here, we perform a principled benchmark of single-cell perturbation prediction that re-examines both model performance and the metrics used to assess it. Rather than introducing yet another architecture, we focus on disentangling genuine predictive signals from artifacts arising from dataset structure, sparsity, and metric design. We benchmark several representative computational models across three chemical perturbation datasets under both out-of-distribution and partially-in-distribution regimes. To contextualize performance, we use dataset-specific reference bounds, as it has been suggested in previous studies^20^, through *CrossSplit* and systematically evaluate the alignment between predicted and observed cellular states at single-cell resolution. We further investigate failure modes of widely used evaluation metrics, identifying scenarios in which they mis-rank models or overestimate performance.

We, in particular, examine commonly-used average-based and distribution-based evaluation metrics using real perturbation datasets, simulations, and controlled noise experiments. We show that average-based metrics are strongly influenced by expression scale and sparsity. Moreover, the Wasserstein distance can yield misleading divergence estimates in high-dimensional settings under variance scaling, and Energy distance may overlook disruptions in gene-gene interactions in certain contexts. We propose a localized Energy distance and a clustering-based *Mixing Index*, which quantifies the co-clustering of predicted and observed perturbed cells. Lastly, we show that a subset of genes, which we term *trivial*, can artificially inflate apparent model performance in differential expression-based evaluations. Together, our results reveal fundamental shortcomings in both existing evaluation practices and current modeling approaches and provide practical guidelines for more reliable benchmarking of single-cell perturbation response models.

## Results

### Overview

To assess the performance of deep-learning models for single-cell perturbation response prediction (Figure 1A), we employed a unified evaluation framework termed CrossSplit, designed to examine both model generalization and the reliability of evaluation metrics, while explicitly clarifying the range of achievable performance.

**Figure 1.**
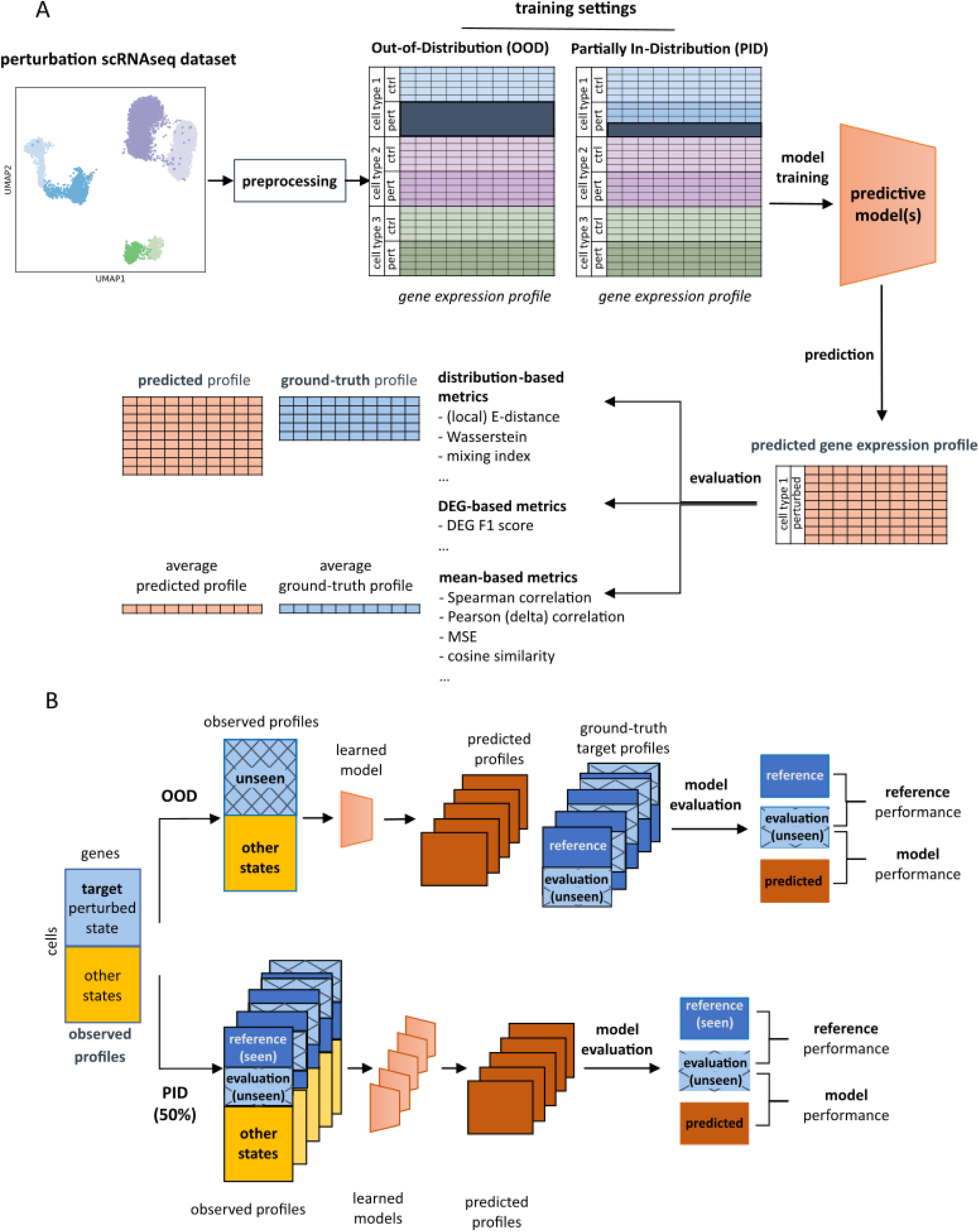
Overview of the training, prediction, and evaluation framework for a single-cell RNAseq perturbation model using CrossSplit. **(A)** The model was trained under two generalization settings. In the OOD setting, all perturbed cells from the target cell type were removed from the training dataset, tasking the model to generate the unseen perturbation-cell type expression profiles. In the PID setting, a specified percentage of the target perturbed cell type was retained during training. After training, the model generated predicted perturbation expression profiles, which were evaluated against ground-truth profiles using both cell-averaged metrics and metrics that consider individual single-cell expression profiles. **(B**) The CrossSplit training and evaluation framework. For the OOD setting, the unseen ground-truth perturbed target cell type was repeatedly split into a reference subset (treated as the best achievable prediction) and an evaluation subset (unseen ground truth). For the PID setting, the target perturbed cell type was split several times into reference and evaluation subsets, with the specified PID percentage determining the proportion assigned to the reference set. The reference subset was then incorporated into the training data used to train the models. In both the OOD and PID settings, evaluation metrics were calculated between (i) the reference subset and the evaluation subset and (ii) the model predictions and the evaluation subset.

We evaluated commonly used metrics, including correlation, Energy distance (E-distance), and Wasserstein distance, alongside newly proposed criteria, with the goal of assessing their effectiveness for benchmarking perturbation-response predictions. These metrics were examined from multiple perspectives, including noise-based analyses and controlled simulation experiments, allowing us to systematically characterize their strengths and limitations.

We considered two complementary generalization settings within CrossSplit (Figure 1B). In the *Out-Of-Distribution* (OOD) setting, all perturbed cells from a target condition are withheld from training and used exclusively for evaluation, thereby testing model performance on entirely unseen perturbations. The withheld cells are randomly divided into a reference group and an evaluation group, and the reference-evaluation comparison serves as a proxy for the upper bound of achievable performance under that condition. In this setting, the reference model simulates the predictions expected from a perfect model. In the *Partially In-Distribution* (PID) setting, a fraction of target perturbed cells (e.g., 50%) is included during training (reference group), while the remaining cells are held out for evaluation. Here, reference-evaluation comparisons instead represent an expected lower bound on achievable performance, as predictive models can leverage both the included target cells and information from other conditions.

In both settings, predictions from all models—including reference, baseline, and complex predictive models—are evaluated against the same held-out evaluation group. Repeating this procedure across multiple random reference-evaluation splits ensures robustness to sampling variability. Using this framework, we evaluated two representative deep-learning models, CPA^21^ and scPRAM^7^, alongside a simple conditional autoencoder (CAE), multiple baseline strategies, and an idealized reference model defining the theoretical performance limit.

The *Results* section is organized around three closely related questions. First, we assess how well existing deep-learning models perform under OOD and PID settings when rigorously evaluated using more reliable metrics. Second, we investigate the behavior and failure modes of commonly used evaluation metrics themselves, using real datasets, controlled simulations and noise experiments. Third, we show that a subset of genes, which we term *trivial genes*, can artificially inflate apparent model performance in differential gene expression analyses. Together, these analyses identify key shortcomings in current evaluation practices and provide a more reliable foundation for guiding the development and benchmarking of perturbation prediction methods.

### CrossSplit Evaluation in OOD Settings Reveals Poor Performance of Existing Deep Learning Methods

Figure 2 and Supplementary Figure 1 present the evaluation results for post-perturbation predictions generated by two complex deep-learning methods, CPA and scPRAM, along with reference and baseline models under the OOD setting across several chemical perturbation datasets. Across all datasets and evaluation criteria, we observe a consistent pattern: complex deep-learning models do not outperform simple models and baselines, and their limitations become most apparent when evaluated using distribution-aware metrics, where they have a substantial performance gap from the optimal reference model.

**Figure 2.**
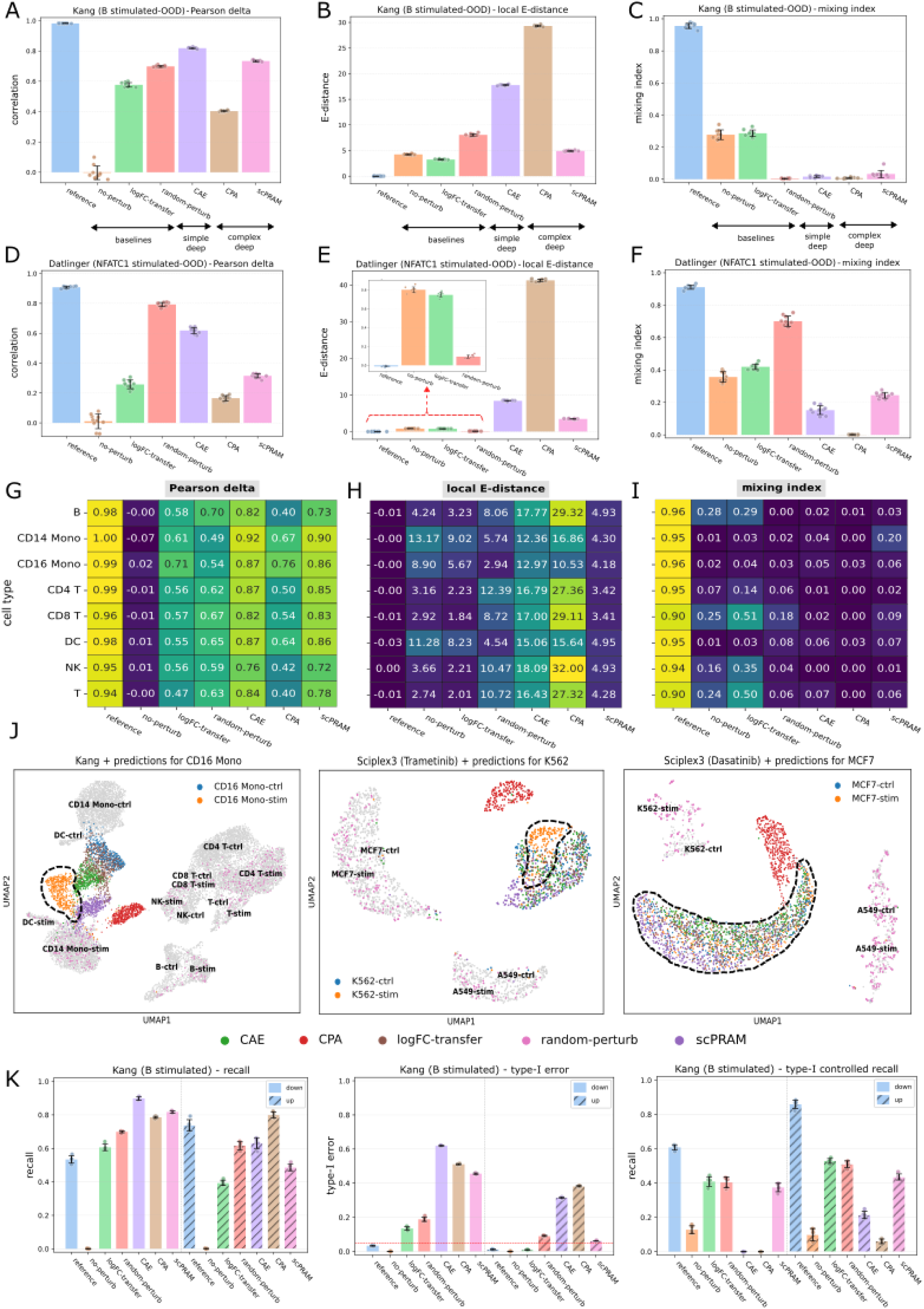
CrossSplit OOD evaluation results. **(A-F)** Bar plots showing Pearson delta, local E-distance and mixing index metrics for evaluating multiple models, including baseline methods (no-perturb, logFC-transfer, and random-perturb), a simple deep learning model (CAE), and two complex deep learning models (CPA and scPRAM), in predicting expression profiles of **(A-C)** the IFN-β stimulated B cells in the Kang dataset and **(D-F)** the CD3 antibody stimulated NFATC1-knockout Jurkat cells in the Datlinger dataset. **(G-I)** Heatmaps of Pearson delta, local E-distance and mixing index metrics across all cell types in the Kang dataset. Each model was trained to predict the stimulated state of each cell type under the OOD setting. Values represent the average performance across CrossSplit repetitions for baseline and deep learning models, as well as the reference model. **(J)** UMAP visualizations of ground-truth control and perturbed single cells (non-target cell types are in grey) alongside predicted single cell profiles (colored) for representative examples: CD16 monocytes under IFN-β stimulation in the Kang dataset (left), K562 cells treated with Trametinib from Sciplex3 (middle), and MCF7 cells treated with Dasatinib from Sciplex3 (right). For UMAP visualizations, scaling and PCA were performed on the whole dataset, comprising ground-truth cells subjected to the same perturbation across different cell types together with unperturbed cells. Model predictions were transformed using the same scaling parameters and projected into the precomputed PCA space, after which UMAP embeddings were computed on the combined dataset including both ground-truth cells and model-predicted profiles. **(K)** Sensitivity analysis of models in identifying true DEGs for stimulated B cells in the Kang dataset. Panels display recall values (left), type-I error (middle), and type-I-controlled recall (right) as explained in the main text, evaluated separately for up- and down-regulated DEGs.

Because many commonly used measures such as correlation on absolute expression levels and Wasserstein distance exhibit some major limitations (analyzed in later sections), we focus here on more reliable metrics, including Pearson delta, local E-distance, and the mixing index.

#### Complex models fail to outperform simple models under OOD generalization

Comparative prediction performance for the perturbed state of cell type B in Kang (Figures 2A-C) and NFATC1-knockout Jurkat cell lines from Datlinger (Figures 2D-F) are shown. The heatmaps summarizing evaluation results across full datasets are provided in Figures 2G-I (Kang) and Supplementary Figure 1 (Datlinger and Sciplex3). Each number in the heatmaps represents the average value of the corresponding metric obtained through the CrossSplit evaluation framework. The heatmap of evaluation results for other less reliable metrics across several datasets are provided in Supplementary Figures 2,3.

As expected, the Pearson delta score for the no-perturb baseline is consistently close to zero. Across the Kang dataset, CPA never outperforms the simple CAE model and scPRAM also fails to surpass CAE for most cell types. Notably in Datlinger, both CPA and scPRAM perform substantially worse than the CAE and random-perturb baselines (Figure 2G and Supplementary Figure 1A).

In Datlinger, Pearson delta scores for the reference model are overall lower than Kang, likely reflecting smaller transcriptional shifts between unperturbed and perturbed cells. This dataset-specific structure limits the dynamic range of achievable scores for this metric and highlights the importance of interpreting model performance relative to dataset-specific upper bounds rather than absolute metric values.

In the Sciplex3 dataset (Supplementary Figure 1D), we observe cases (e.g., MCF7 cells treated with Sorafenib) where the Pearson delta of the reference model is far from optimal (below 0.3). This observation suggests that this metric may be unreliable in such cases or may be more appropriate when applied to carefully selected gene subsets.

The limitations of complex models become more pronounced when evaluated using distribution-based metrics. Across all datasets, CPA consistently underperforms most baselines in both local E-distance and mixing index, indicating poor recovery of the perturbed cell-state distribution. scPRAM performs modestly better than CPA on these metrics but remains substantially worse than the reference model, especially for the mixing index, where a large performance gap persists (Figures 2H,I; Supplementary Figures 1B,C,E,F).

#### CrossSplit baselines reveal dataset structure

A detailed examination of the CrossSplit results for the reference, no-perturb and random-perturb models might provide insight into the intrinsic structure of each dataset. In the Sciplex3 dataset, there are cases in which the no-perturb baseline achieves a high mixing index close to the reference score, indicating that perturbed and unperturbed cells occupy nearby regions of expression space (Supplementary Figure 1F). Likewise, the elevated mixing index observed for several cases of the random-perturb in Datlinger, relative to Kang, suggests closer proximity among perturbed states across different cells (Supplementary Figure 1C). This behavior aligns with expectations, since Kang comprises multiple peripheral blood mononuclear (PBMC) cell types, whereas Datlinger consists of a single cell line (Jurkat) subjected to different gene knockouts, which will go under stimulation.

It is important to note that an evaluation metric may not be equally appropriate across all datasets. Beyond accurately assessing model performance, an informative metric should yield a clear performance gap between the reference model and the baselines. When such a gap is absent, this may reflect that the perturbation effect is confined to a relatively small subset of genes.

#### Embedding visualizations corroborate distributional failures

UMAP embeddings are presented in Figure 2J, including one example from Kang (left, CD16 monocytes) and two from Sciplex3 (middle and right, K562 treated with Trametinib and MCF7 treated with Dasatinib, respectively). In all cases, CPA predictions fail to recapitulate the geometry of the true perturbed cell distributions. Noticeable deviations are also evident for scPRAM as well as other models and baselines in the first two example UMAPs, underscoring the intrinsic difficulty of the task.

#### Differential expression-based evaluations can be misleading

Finally, we examined model performance using differential gene expression (DEG) analysis. Evaluating the recovery of DEGs is often considered more biologically meaningful. To select an appropriate statistical approach for computing differential expression p-values between perturbed and unperturbed cells, we employed our single-cell RNA-seq simulator. Details of this selection procedure are provided in the Supplementary Notes 2,3 and Supplementary Figure 10. Using the selected approach, we assessed the sensitivity of different models in identifying true DEGs, defined as genes with p-values below 0.05, in the Kang dataset (Figure 2K; Supplementary Figure 4). Because our primary goal was to construct a ground-truth reference for comparative evaluation across models, we did not apply multiple-testing correction.

Notably, the recall of the reference model was well below one, largely due to the high sparsity of single-cell data, and was lower than that of most other models, especially for down-regulated genes (Figure 2K, left). This apparent underperformance is driven by high values of type-I error rates across most models (Figure 2K, middle), which renders recall estimates unreliable. After adjusting the p-value threshold for DEG detection to control the type-I error rate below 0.05, recall for complex models, especially CPA, dropped below that of most baselines (Figure 2K, right). These results demonstrate that DEG recovery-based evaluations can substantially overestimate model performance when error rates are not properly controlled, underscoring the need for cautious interpretation of such metrics.

### Complex Models Fail to Reconstruct Target Distributions of Perturbed Cells under Standard Train-Test Split Settings

To further examine model performance beyond strict OOD generalization, we evaluated the effect of partial inclusion of target perturbed cells during training (PID setting). Specifically, models were trained with varying fractions of cells from the target perturbed condition included in the training set. As the PID percentage increases, models receive progressively more direct information about the target condition, theoretically reducing the prediction task difficulty. Under this regime, complex models would be expected to outperform the reference model because, in addition to using the same information as the reference model, they can incorporate data from other perturbed cells during training.

Despite partial exposure to the target condition, the mean-expression prediction performance of both complex models, CPA and scPRAM, consistently underperformed the CAE and the reference model, across most PID percentages. Nevertheless, across PID levels, we observed slight improvements for CPA, reflected by small gains in Pearson delta (Figures 3A-B, left).

**Figure 3.**
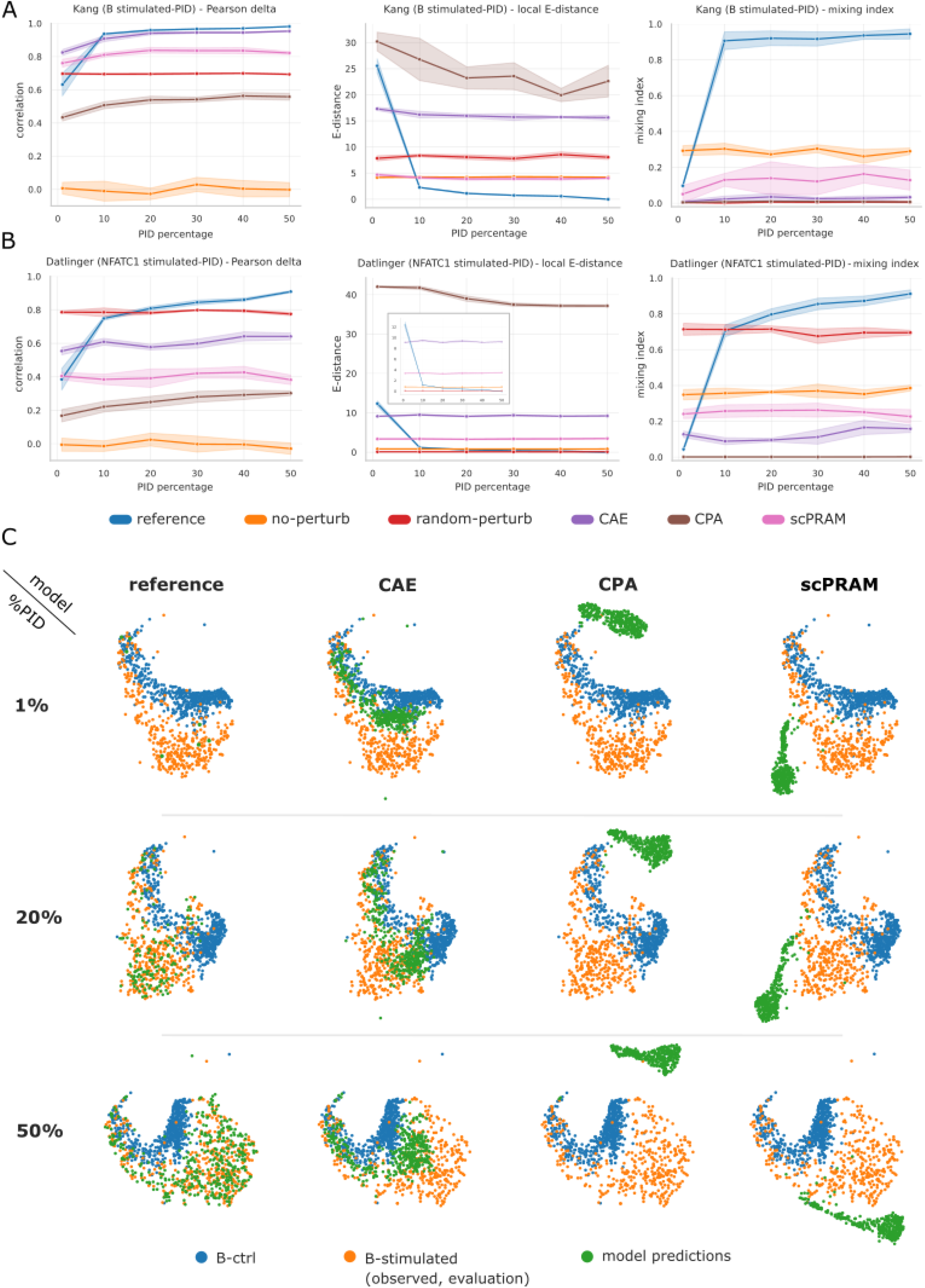
Model performance under the PID training setting assessed using CrossSplit evaluation. **(A-B)** Pearson delta, local E-distance and mixing index for models trained with increasing proportions (1%, 10%, 20%, 30%, 40%, 50%) of perturbed target cell types included during training. For each condition, performance represents the mean across five independent CrossSplit repetitions, with shaded regions denoting the standard deviation. **(A)** Performance on predicting the stimulated state of B cells in the Kang dataset. **(B)** Performance on predicting the stimulated state of NFACT1-knockout Jurkat cells in the Datlinger dataset. **(C)** UMAP visualizations of ground truth B cells from the Kang dataset alongside model predictions under PID training settings in which 1%, 20% or 50% of stimulated B cells were included in the training data. Model-predicted cells are shown in green. The evaluation subset of stimulated B cells (unseen set) is shown in orange and unperturbed (control) B cells are displayed in blue. Scaling and PCA were performed on the whole dataset, comprising ground-truth cells subjected to the same perturbation across different cell types together with unperturbed cells. Model predictions were transformed using the same scaling parameters and projected into the precomputed PCA space. For each PID percentage, UMAP embeddings were computed on the combined dataset consisting of ground-truth cells (including the observed evaluation subset) and all model-predicted profiles corresponding to that PID level.

Distribution-based measures provide a more stringent assessment of model behavior. Both local E-distance and mixing index indicate that complex models fail to fully reconstruct the distribution of target perturbed cells, even when trained with substantial fractions of those cells (Figures 3A-B, middle and right). While minor improvements were observed for CPA in local E-distance and for scPRAM in the mixing index for specific cell types and conditions, predicted cell distributions remained markedly divergent from the true perturbed distributions, as illustrated in the example UMAP visualizations (Figure 3C). We further repeated the evaluation using varying sample sizes which avoided the up-sampling of the reference group at lower PID levels. This analysis again confirmed the poor performance of complex models under the PID setting (see Supplementary Note 4 and Supplementary Figure 5).

### Average Expression-Based Metrics Are Misleading When Applied to Absolute Expression Values

A common strategy for comparing predicted and observed single-cell expression profiles is to aggregate gene expression across cells and compare vectors of mean expression, either across genes or on a per-gene basis across cell types (Figure 4A). Under CrossSplit evaluation in the OOD setting, however, average-based metrics such as across-genes mean squared error (MSE) and cosine similarity failed to consistently distinguish the reference model from baselines (Supplementary Figures 2,3), particularly in the Datlinger and Sciples3 datasets, indicating their limited discriminative power.

**Figure 4:**
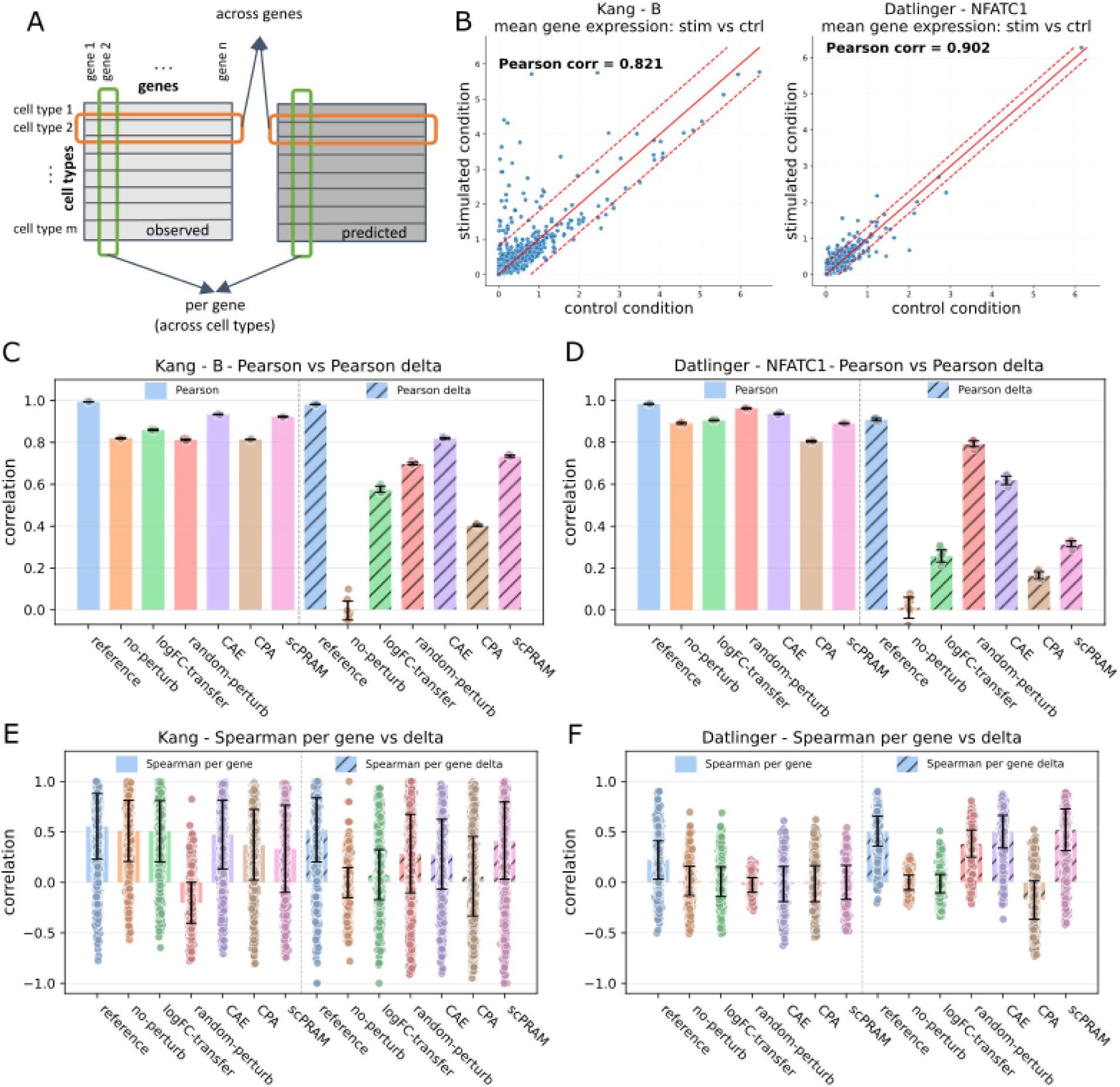
Evaluation of correlation-based metrics. **(A)** Schematic illustrating model performance assessment using correlation-based metrics. Each row in the observed (ground truth) and predicted matrices represents the average expression profile of a perturbed cell type across single cells, and each column represents a gene. Across-genes evaluation refers to applying a correlation metric between the observed and predicted average profiles of the same cell type (orange box). In contrast, per-gene or across-cell-types evaluation applies the correlation metric to the averaged profiles of a single gene across different cell types from observed and predicted data (green box). **(B)** Scatter plots of mean expression values of individual genes for unperturbed vs perturbed cells with corresponding across-genes Pearson correlation values for stimulated B cells from the Kang dataset (left) and stimulated NFATC1-knockout cells from the Datlinger dataset (right). In each panel, the two dashed lines represent *y* = *x* ± *b*, where *b* is chosen such that 99% of the genes lie within the bounded area between these lines. **(C-D)** Bar plots comparing across-genes Pearson (solid) and Pearson delta (hatched) correlations for model performance averaged over 10 CrossSplit repetitions: **(C)** stimulated B cells in the Kang dataset and **(D)** stimulated NFATC1-knockout Jurkat cells in the Datlinger dataset. **(E-F)** Bar plots showing per-gene Spearman correlation (solid) and its delta-corrected variant (hatched) across perturbed cell types for **(E)** Kang and **(F)** Datlinger datasets.

Similarly, CrossSplit evaluation using across-genes Spearman correlation (Supplementary Figures 2,3) revealed that in many cases, the theoretically perfect reference model underperformed several baselines and benchmark models (Supplementary Figures 6B,D; values rescaled relative to the reference). This observation highlights the unreliability of across-genes Spearman correlation as a performance metric in this context.

Comparing the mean expression values of individual genes between unperturbed and perturbed cells (Figure 4B) showed that the majority of genes exhibit comparable average expression across control and perturbed conditions, resulting in high across-genes Pearson correlations for the no-perturb baseline. Thus, across-genes Pearson correlation on absolute expression values are largely driven by expression scale rather than perturbation-induced changes.

Applying Pearson correlation to expression changes instead of absolute values mitigates this scale-driven behavior (Figures 4C,D). This delta-corrected metric, referred to as Pearson delta (Pearson_Δ_), provides a more biologically meaningful assessment by evaluating whether the direction of gene expression change is correctly predicted, as used in previous studies^5,9,18^. Notably, under Pearson delta, complex deep-learning models (CPA and scPRAM) underperformed the simple CAE model in both the Kang and Datlinger datasets and fell below the random-perturb baseline in Datlinger (Figures 4C,D).

In contrast, the analogous delta-corrected Spearman correlation (Spearman delta) showed inconsistent behavior: in the Datlinger dataset, the reference model was again outperformed by the CAE in several cases (Supplementary Figures 6F), suggesting that Spearman delta may not be reliable, at least in certain experimental contexts. Per-gene Spearman correlation evaluated across cell types exhibited limited discriminatory power in Kang (Figure 4E), and its delta-corrected counterpart performed similarly poorly in Datlinger (Figure 4F). Rescaling per-gene scores relative to the reference model for individual genes further confirmed that, in many cases and for many genes, the reference model is outperformed by other models under either formulation (Supplementary Figures 6G,H).

Although average expression-based metrics are computationally efficient and intuitively interpretable, they obviously fail to capture the pronounced cell-to-cell heterogeneity inherent in single-cell data and do not reflect gene-gene dependencies. These limitations motivate the use of distribution-based evaluation metrics that explicitly assess how well a perturbation prediction model reconstructs the full distribution of perturbed cellular states. Accordingly, we evaluated distributional fidelity of models using two widely adopted distribution-based measures, the Wasserstein distance and the Energy distance (E-distance), together with a clustering-based metric, the Mixing Index.

### Wasserstein Distance Can Exhibit Fundamental Failure Modes in Medium- and High-Dimensional Gene Expression Data

CrossSplit evaluation results for the Wasserstein distance in the Kang and Datlinger datasets, rescaled relative to the reference model by subtracting the reference score (Figure 5A), revealed striking inconsistencies. In several cases, baseline and benchmark models achieved negative rescaled values, indicating apparently better performance than the theoretically perfect reference model. This counterintuitive outcome suggests that the Wasserstein distance can yield misleading divergence estimates in this setting.

**Figure 5.**
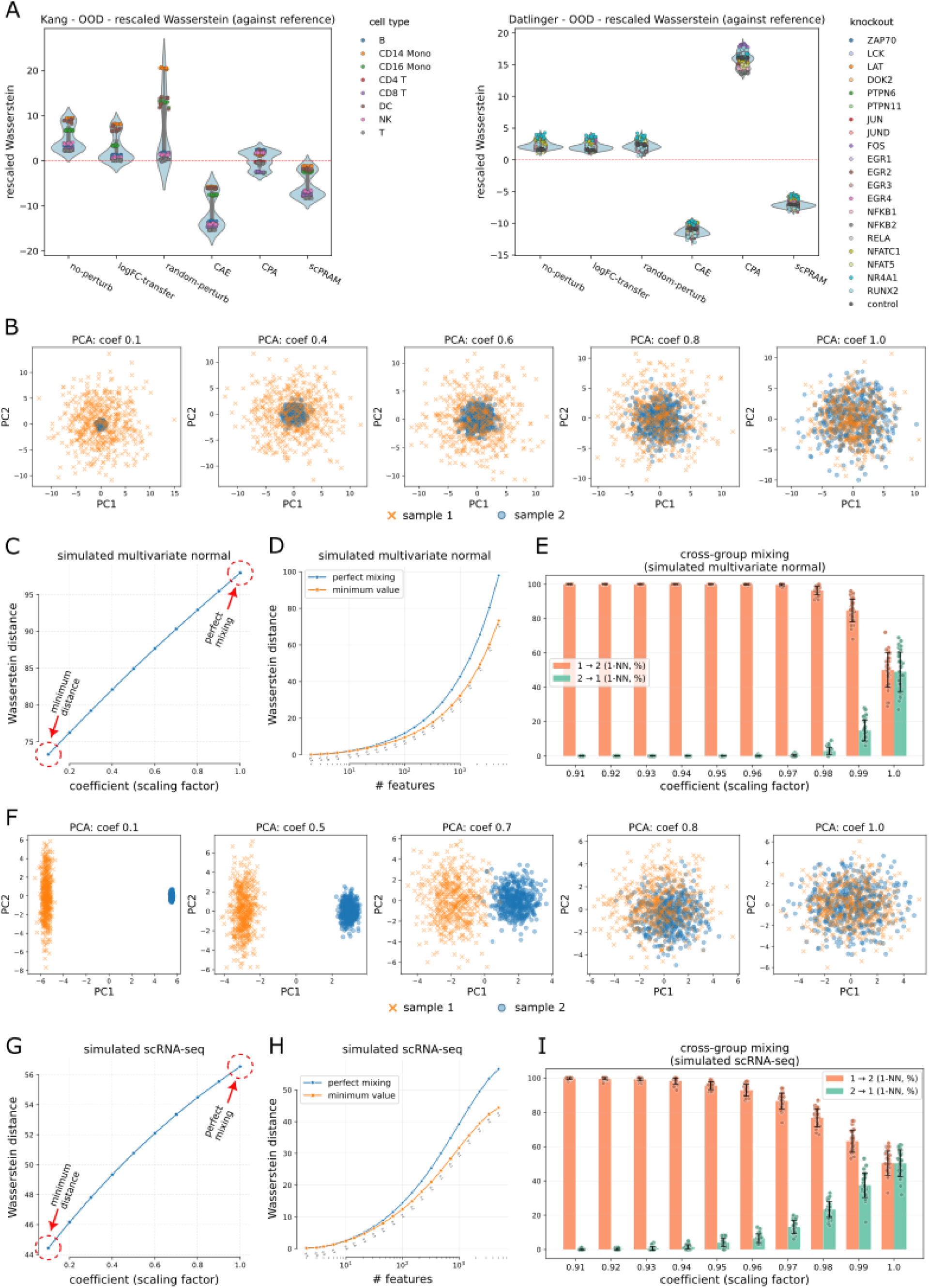
Evaluation of the Wasserstein distance metric on simulated data. **(A)** CrossSplit evaluation of model performance using the Wasserstein distance, rescaled relative to the reference model, for predicting the stimulated state of each cell type in the OOD setting for the Kang (left) and Datlinger (right) datasets. Values below zero indicate cases in which a model outperforms the reference model in the same evaluation repeat. **(B-E)** Experiments using Gaussian simulations: **(B)** PCA visualizations of two simulated samples of 500 cells, each drawn from a 5,000-dimensional multivariate normal distribution with zero mean and identity covariance. From right to left, the variance of the blue sample progressively decreases. **(C)** Wasserstein distance between the two samples in panel B plotted against the variance-scaling coefficient controlling the variance of the blue sample. **(D)** Wasserstein distances between the two samples in panel B as a function of the number of features. The blue curve corresponds to samples with equal variance (coefficient = 1), while the orange curve shows the minimum Wasserstein distance among the explored variance-scaling coefficients at each dimensionality. The coefficient at which this minimum occurs is indicated below the orange curve for selected dimensionalities. **(E)** Nearest-neighbor relationships between two simulated samples drawn from 5,000-dimensional multivariate normal distributions (100 cells each). Sample 1 has identity covariance (scale = 1.0), whereas the covariance of sample 2 is scaled from 1.0 down to 0.91. The x-axis denotes the variance-scaling coefficient of sample 2. For each setting, *i→j (1-NN, %)* indicates the percentage of cells in sample *i* whose nearest neighbor belongs to the other sample *j*. **(F-I)** Analogous experiments on samples generated using our single-cell RNA-seq simulator with negative binomial counts. Panels **F-I** mirror panels **B-E**, respectively, with log-normalized expression values: **(F)** PCA visualizations of two simulated samples (500 cells each). From right to left, the read-count variance of the blue sample progressively decreases. (**G)** Wasserstein distances plotted against the coefficient controlling the variance of the blue sample. **(H)** Wasserstein distances as a function of the number of features, with color coding analogous to panel D. **(I)** Nearest-neighbor relationships.

To further investigate the behavior of distribution-based metrics, we conducted controlled simulations with known ground-truth distributions. We first generated two samples from a 5,000-dimensional multivariate normal distribution with zero mean and identity covariance. Initially, the samples were well mixed as shown in the PCA space (Figure 5B, coefficient = 1). We then progressively reduced the variance of one sample (sample 2; blue circles), causing it to contract toward the center of the distribution (Figure 5B, coefficients 1 → 0.1). Conceptually, this shrinkage should increase divergence between the two samples.

Consistent with this expectation, E-distance increased as the variance of one sample decreased (Supplementary Figure 7A). In contrast, the Wasserstein distance paradoxically decreased (Figure 5C) and, surprisingly, failed to assign the minimum distance to the perfectly mixed sample (coefficient = 1). To study the dependence of this phenomenon on dimensionality, we repeated this simulation across varying numbers of features. As shown in Figure 5D, the variance-scaling coefficient minimizing the Wasserstein distance shifted from 1 toward smaller values as dimensionality increased. Thus, in higher-dimensional spaces, a more concentrated distribution can appear *closer* to the original distribution than one drawn from the same underlying parameters, demonstrating a fundamental failure mode of the Wasserstein distance.

To explain the counterintuitive behavior of the Wasserstein distance under variance scaling, we performed a nearest-neighbor analysis on high-dimensional multivariate normal simulated data, revealing a pronounced asymmetry whereby cells from a more diffuse distribution preferentially match to a more compact one (Figure 5E). This Euclidean nearest-neighbor asymmetry provides an intuitive explanation for why Wasserstein distance can decrease as one distribution becomes increasingly concentrated in high-dimensional spaces (see Supplementary Note 5 for more details)

We repeated these analyses using our single-cell RNA-seq simulator to better reflect realistic data properties (Supplementary Note 1). Read counts were sampled from a negative binomial distribution, normalized to a total of 10⁴ counts per cell, and subsequently log-transformed. We generated two identically distributed samples with a mean of 100 and number of successes *r* = 1 across 5,000 genes to simulate the count matrices. We then progressively reduced the variance of sample 2 (Figure 5F, PCA plots of log-normalized expression values).

Again, Wasserstein distance decreased as the variance of one sample was reduced (Figure 5G), contrary to the expected divergence. In contrast, both E-distance and the mixing index behaved as anticipated: E-distance increased and mixing index decreased (Supplementary Figures 7B,C). Additional simulations across varying dimensionalities and nearest-neighbor analyses (Figures 5H,I) reproduced similar patterns observed in the Gaussian simulations.

Taken together, these results demonstrate that in medium- and high-dimensional settings, characteristic of single-cell transcriptomic data, a lower Wasserstein distance does not necessarily indicate greater similarity between distributions. This raises serious concerns regarding the use of Wasserstein distance as a primary metric for evaluating perturbation prediction models in single-cell RNA-seq and underscores the need for more robust, structure-aware distributional metrics.

### E-distance Might Fail to Capture Disruptions in Gene-Gene Interactions

To further evaluate the behavior of distribution-based metrics, we designed a controlled shuffling-noise experiment (Figure 6A; see *Methods, Noise sensitivity analysis*), which preserved per-gene marginal distributions but disrupted potential gene-gene dependencies within the noisy split. Our goal was to measure the divergence between each noisy split and the unchanged test split using distribution-based evaluation metrics and assess the sensitivity of different metrics to perturbations in gene-gene interactions.

**Figure 6.**
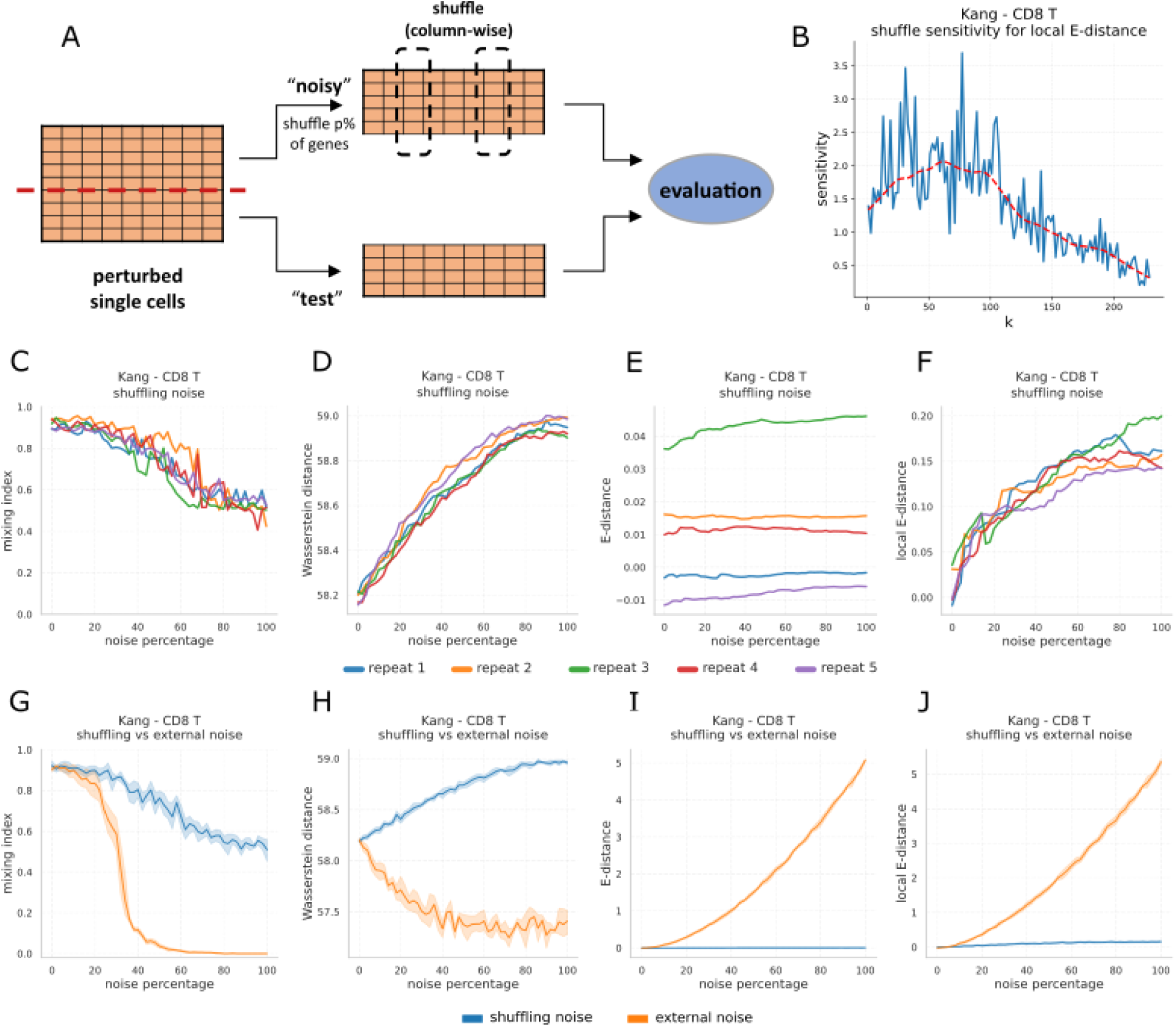
Performance of distribution-based metrics in capturing gene-gene interactions and matrix-level noise. **(A)** Schematic of the controlled shuffling-noise experiment. True perturbed single cells of the target cell type are randomly split into a *test* set and a *noisy* set. In the noisy set, a given percentage of genes has their expression values randomly shuffled across cells, after which divergence between the noisy and test sets is quantified. The procedure is repeated across multiple random splits. **(B-F)** Analyses regarding (nested) shuffling noise experiment. **(B)** Sensitivity of local E-distance to shuffling-induced disruptions across different neighborhood sizes *k* in CD8 T cells from the Kang dataset. Values of **(C)** mixing index, **(D)** Wasserstein distance, **(E)** E-distance and **(F)** local E-distance, computed between the noisy and test sets of stimulated CD8 T cells in the Kang dataset across noise levels ranging from 0% to 100%. **(G-J)** Comparing the effect of non-nested shuffling-noise and matrix-level noise-addition (external noise) on stimulated CD8 T cells from the Kang dataset, measured by **(G)** mixing Index, **(H)** Wasserstein distance, **(I)** E-distance and **(J)** local E-distance. Shaded regions denote variability across five repetitions.

As the proportion of shuffled genes increased, both the mixing index and the Wasserstein distance between the noisy and test splits deteriorated (Figures 6C,D for Kang; Supplementary Figures 8A,B for Datlinger), indicating sensitivity to disruptions in gene-gene relationships. In contrast, E-distance exhibited a notable limitation: in certain datasets or cell types, the variability of E-distance between noisy and test splits due to sampling randomness exceeded the changes induced by shuffling. This issue was particularly evident for certain cell types in the Kang dataset (e.g., CD8 T cells; Figure 6E), but was not observed in Datlinger (Supplementary Figure 8C). These results suggest that while E-distance captures global distributional shifts, it may lack sufficient sensitivity to detect more subtle disruptions of gene-gene interactions in certain contexts.

To address this limitation, we introduced *Local E-distance*, in which pairwise distances are computed only within local neighborhoods rather than across all cell pairs. This modification greatly enhanced sensitivity to shuffling-induced disruptions relative to sampling variability (Figure 6F; Supplementary Figure 8D). The neighborhood size *k* influences the performance of local E-distance; we therefore selected *k* by maximizing a predefined *sensitivity* criterion (see *Methods*), with a lower bound of *k* = 30 to avoid unstable and overly local estimates. Example sensitivity curves for Kang CD8 T cells and Datlinger NFATC1-knockout cells are shown in Figure 6B and Supplementary Figure 8E, respectively.

To complement the shuffling-noise analysis, we performed a second noise-addition experiment that introduced external noise and perturbed the expression matrix at the level of individual entries. In the Kang dataset, the distance of noisy and test splits based on both mixing index and (local) E-distance steadily increased with noise level, with greater intensity than observed under shuffling-noise experiment (Figures 6G,I-J). By contrast, Wasserstein distance failed to consistently increase with noise level, indicating an inability to capture distributional divergence as the noisy split departed from the test split (Figure 6H). Again, this provides additional evidence that questions the suitability of Wasserstein distance as a divergence metric in this context. However, this misbehavior was not observed in the Datlinger dataset (Supplementary Figure 8G), suggesting that dataset-specific characteristics may modulate the Wasserstein’s failure modes.

### Identification of top-ranking differentially expressed genes is artificially easy due to the inflated prevalence of zeros in gene expression data

Many studies developing single-cell perturbation prediction models evaluate their performance on a subset of top-ranked differentially expressed genes (DEGs), motivated by their presumed relevance to transcriptional responses. However, a well-performing perturbation prediction model should accurately predict expression values beyond this narrow subset as well, including lower-ranked DEGs and genes that are not significantly differentially expressed. Focusing exclusively on top-ranked DEGs risks providing an incomplete and potentially misleading assessment of model performance.

To examine the impact of gene selection, we categorized genes into three groups based on sparsity and differential expression significance: *trivial*, *non-trivial* and *non-significant* (see details in *Methods*). Trivial DEGs are characterized by extreme sparsity, such that minimal predictions, i.e., assigning small non-zero expression values to a limited subset of the perturbed cells for the corresponding gene, can suffice for a model to correctly predict the gene as a DEG. Two representative examples from Kang B cells are shown in Figures 7A,B. Because these genes are easy to predict as DEG, evaluations restricted to trivial DEGs can artificially inflate performance estimates in differential expression analyses.

**Figure 7.**
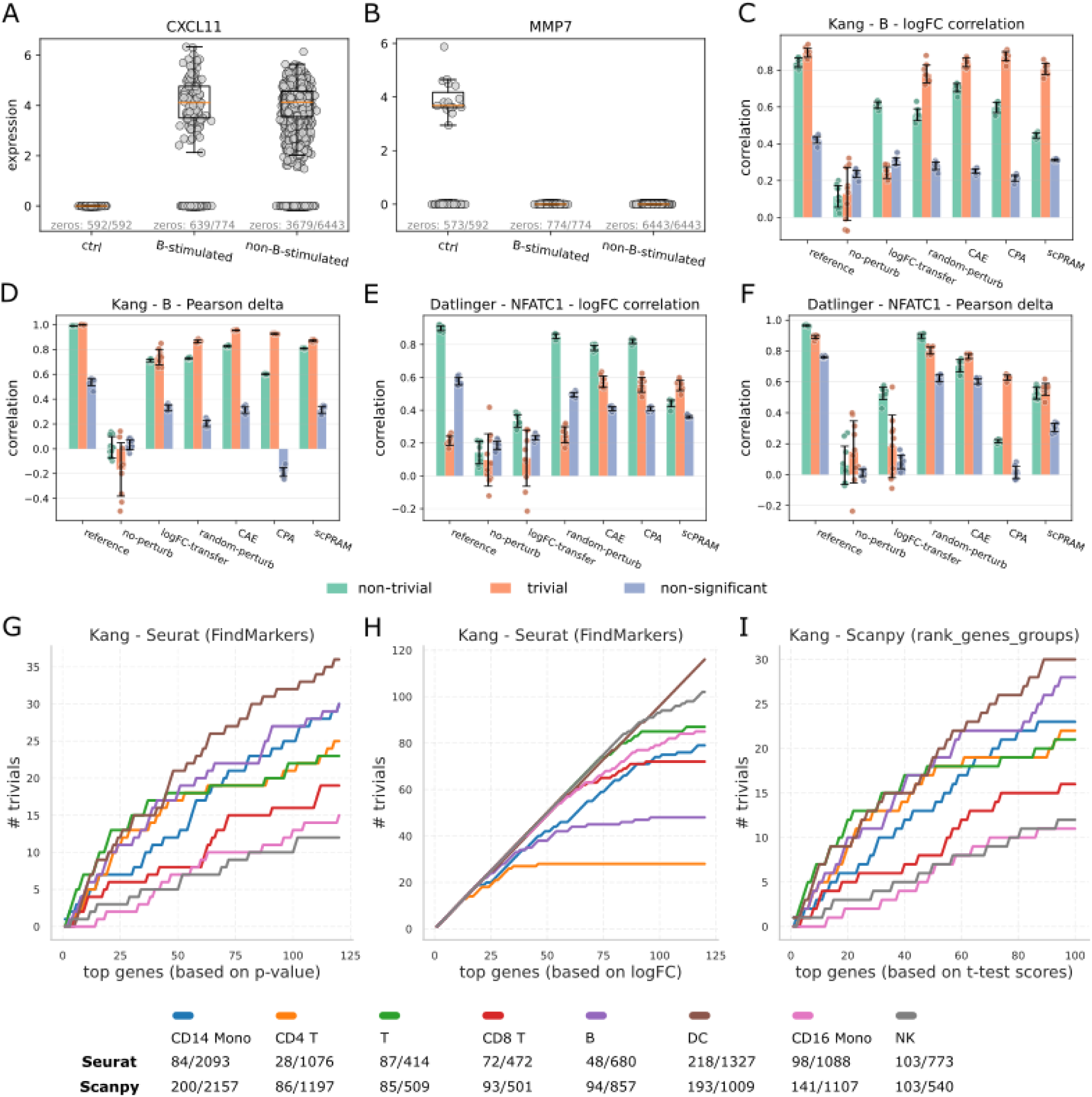
CrossSplit evaluation of model performance across gene categories. **(A,B)** Two representative trivial genes in stimulated B cells from the Kang dataset: **(A)** *CXCL11* and **(B)** *MMP7*, corresponding to DEGs with the highest and lowest log fold-change (logFC), respectively. For each gene, dot plots display the distribution of expression values across all single cells in the population of control cells (‘ctrl’), target perturbed cells (‘B-stimulated’) and non-target cell types subjected to the same perturbation (‘non-B-stimulated’). Boxplots summarizing non-zero expression values for each gene and population are overlaid. **(C)** logFC correlations and **(D)** Pearson delta values from CrossSplit evaluation of models on trivial, non-trivial and non-significant genes in stimulated B cells from Kang. **(E)** logFC correlations and **(F)** Pearson delta values from CrossSplit evaluation on the same gene categories for stimulated NFATC1-knockout cells from Datlinger. **(G-I)** Prevalence of trivial genes among top-ranked DEGs across all eight cell types in the Kang dataset, ranked by **(G)** p-value, **(H)** logFC among significant genes (p-value ≤ 0.05) obtained using Seurat’s *FindMarkers* function, and **(I)** absolute z-scores computed using Scanpy’s *rank_genes_groups* function with a *t-test*.

Metrics such as logFC correlation and Pearson delta, which assess expression changes relative to control cells, are particularly susceptible to this bias. CrossSplit evaluation results across gene subsets (Figures 7C,D for Kang B cells, Figures 7E,F for Datlinger NFATC1-knockout cells) showed substantial variation in model performance depending on gene category. Notably, CPA and scPRAM achieved markedly higher scores on trivial DEGs than on the non-trivial group in Kang B cells. On the other hand, quantifying the prevalence of trivial DEGs among top-ranked genes revealed that a substantial proportion of top-ranked DEGs, frequently used by a variety of studies for model benchmarking, are in fact trivial across many cell types (Figures 7G-I; Supplementary Figures 9A-C). Together, these results indicate that evaluations confined to top-ranked DEGs can bias performance estimates in control-referenced average-based metrics, underscoring the importance of evaluating models on non-trivial DEGs as well.

In Datlinger NFATC1-knockout cells, logFC correlation further revealed notable inferior performance of the reference model on trivial DEGs compared with the random-perturb baseline and deep-learning models (Figure 7E), highlighting the unreliability of this metric when applied to sparsity-dominated gene subsets. For non-significant genes, the reference model exhibited lower correlations in Kang (Figures 7C,D), consistent with weak and noisy expression differences between control and perturbed cells. However, elevated reference scores for non-significant genes in Datlinger (Figures 7E,F) suggest residual differential expression signals, reflecting inherent difficulty of defining truly non-significant sets in sparse single-cell data.

CrossSplit evaluation results using distribution-based metrics across gene subsets (Supplementary Figures 9D-I) showed that the reference model achieved near-optimal performance under E-distance and the mixing index across all gene categories. In contrast, CPA exhibited a pronounced performance gap relative to the reference model on non-trivial and non-significant genes based on both metrics. While scPRAM showed a relatively higher mixing index on trivial genes in Kang B cells, its performance remained poor on non-trivial and non-significant genes.

In line with earlier findings, Wasserstein distance again demonstrated marked limitations (Supplementary Figures 9F,I), as the reference model failed to outperform other models across any of the gene groups under this metric.

## Discussion

The rapid growth of single-cell perturbation datasets and advances in deep-learning architectures has created an expectation that accurate, generalizable prediction of cellular responses is within reach. Our results challenge this assumption. Across multiple chemical perturbation datasets, generalization regimes, and evaluation metrics, we find that perturbation prediction models remain far from reconstructing the true distribution of perturbed cellular states. Importantly, this gap persists even when models are partially exposed to the target perturbed cells during training, suggesting that the limitation is not simply a lack of data or model capacity.

A central insight from this study is that progress in perturbation modeling has outpaced the development of reliable evaluation practices. Our findings show that several widely adopted metrics are sensitive to scale, sparsity, and dimensionality, leading to systematic mis-ranking of models or in some cases, assigning better scores to degenerate or trivial predictions than to genuinely more accurate ones in realistic single-cell settings. Consequently, models may appear to perform well while failing to capture cell-to-cell variability and gene-gene dependencies.

Establishing achievable performance bounds is essential in computational biology. Community benchmarks efforts such as the Critical Assessment of Structure Prediction (CASP) illustrate this principle, using experimentally determined structures to define concrete reference standards that guide methodological progress. In this study, we similarly define dataset-specific performance bounds through CrossSplit. In some settings, simple baselines match or even exceed reference-level performance under certain metrics, indicating that those metrics are not well suited for the task. In some datasets, reference-level performance is approached because perturbation effects are weak or restricted to a small subset of genes, making the difference between unperturbed and perturbed responses small and allowing even simple models to achieve performance similar to the reference level. In both cases, strong performance claims should be interpreted with caution, as they may reflect properties of the dataset or metric rather than genuine predictive ability.

Our analyses reveal a mismatch between signals that are easy to predict and those that are biologically informative, whose accurate recovery more faithfully reflects genuine model capability. Differential-expression-based evaluations, particularly when restricted to top-ranked genes, are often dominated by trivial genes arising from zero inflation and sparsity. Success on these subsets does not imply accurate modeling of broader transcriptional responses or cell-to-cell variability. More generally, evaluations that focus on marginal gene-level effects overlook gene-gene dependencies, which are essential for understanding cellular states under different perturbations.

The mixing index and localized E-distance demonstrate how evaluation can be aligned with the structure of single-cell data. By clustering predicted and observed perturbed cells of the target condition together with other cell types and conditions, the mixing index assesses whether predictions align specifically with the target perturbed population rather than unrelated states. This approach is conceptually related to *centroid accuracy* proposed by *Systema*^18^, but operates at the single-cell level. It should be noted that clustering single-cell RNA-seq data is inherently challenging due to high sparsity. We observed substantial differences between Louvain clustering results obtained using Seurat and Scanpy, consistent with prior reports^22^. While sensitive to clustering parameters (e.g., resolution), methodological choices and sample size, the mixing index provides a bounded and interpretable measure of distributional alignment that complements distance-based metrics.

Shuffling-noise experiments demonstrated that global E-distance may lack sensitivity to disruptions in gene-gene dependencies, while its localized variant improves sensitivity at the cost of introducing an influential hyperparameter (i.e., neighborhood size). Simulations further revealed fundamental limitations of the Wasserstein distance in medium- and high-dimensional settings under variance scaling, raising concerns about its reliability for evaluating high-dimensional single-cell expression data, despite its central role in optimal transport-based modeling.

Together, these results suggest that the main barrier to predictive virtual-cell models is not architectural complexity, but the lack of rigorous and context-aware evaluation standards. Careful selection of evaluation metrics and gene subsets is critical, and continued development of robust metrics remains an important open challenge for the field.

## Methods

### Single cell perturbation datasets

***Kang et al.*** *(2018):* The dataset consists of single-cell gene expression profiles from PBMC samples of eight individuals, which were either stimulated with interferon beta (IFN-β) or left untreated. We used preprocessed data adopted from Lotfollahi *et al.*^21^ consisting of cells from 8 distinct cell types and featuring 5,000 highly variable genes (HVGs). The original data is available at NCBI Gene Expression Omnibus (GEO) under accession number GSE96583^23^.

***Datlinger et al.*** *(2021):* Jurkat single-cell gene expression profiles were generated after CRISPR-based targeting of 20 T-cell receptor pathway genes, followed by stimulation of knockout and control cells with an anti-human CD3 antibody. We obtained count data from NCBI GEO under accession number GSE168620^24^. Data preprocessing was performed using Scanpy by filtering cells with at least 100 counts, retaining genes expressed in a minimum of five cells and selecting 5,000 HVGs.

***Srivatsan et al.*** *(2020):* The Sciplex3 dataset contains three cancer cell lines (A549, MCF7, K562) treated with 188 small-molecule compounds at four concentrations, alongside vehicle controls. Count data are available in NCBI GEO under accession number GSE139944^25^. Preprocessing in Scanpy involved filtering cells with at least 100 total counts, retaining genes expressed in at least five cells and selecting 5,000 HVG. For model evaluation, we focused on perturbation response prediction for five representative compounds including Trametinib, Panobinostat, Dasatinib, Sorafenib tosylate and Crizotinib, at a concentration of 1,000 nM.

### Model training settings

We train models under two generalization settings (Figure 1A): In the Out-of-Distribution (OOD), training data include perturbed and unperturbed single cells from all cell types except the target cell type, for which the model has access only to the unperturbed state, while the corresponding perturbed condition is entirely excluded from training. At test time, the model is provided with the unperturbed cells of the target cell type and tasked with predicting their perturbed state. In Partially In-Distribution (PID), a fraction of perturbed cells from the target cell type (e.g., 20%) is included in the training data, while the remaining perturbed cells are held out. Similar to the OOD setting, the trained model is provided with the unperturbed cells of the target cell type to predict their perturbed profiles.

### Evaluation framework

We established a robust framework to evaluate model performance in predicting gene expression profiles (Figure 1B). For each OOD condition, target perturbed cells withheld from training were randomly divided into two groups: a reference group and an evaluation (unseen) group. The reference group served as a proxy for an ideal model which represents the most accurate gene expression profiles achievable under the given condition. Model predictions were compared with the unseen cells (evaluation group), with the number of predicted and observed cells matched. To estimate the upper bound of achievable performance, the same metrics were computed between the reference and unseen groups. This procedure was repeated several times with different random splits to ensure robustness.

In the PID setting, five independent models were trained for each inclusion percentage. For each model, the specified proportion of target perturbed cells was randomly selected as the reference group and included in training, while the remaining cells were withheld as unseen data. Predictions were then evaluated against the unseen cells, with reference-unseen comparisons instead serving as an estimate of the performance lower bound. The reason is that at this time, models can take advantage of the information existing in the included fraction of target cells in addition to the other conditions.

The *comparison size* is considered as half the number of the real target perturbed cells. During evaluation at each PID percentage, the evaluation group is down-sampled to the size of comparison size before comparison. Similarly, the generative models were required to produce the same number of predictions.

### Evaluation metrics

In the following, we summarize commonly used evaluation metrics and introduce our proposed modifications designed to address some of the current limitations.

#### Pearson delta

Correlation-based metrics such as Spearman correlation, Pearson correlation, and the coefficient of determination (R²) are widely used in the previous studies^4,5,9,21^. Typically, correlation metrics are applied to predicted and observed expression profiles after averaging them across cells within each perturbation condition.

To reduce the dominance of correlations by highly expressed genes and to make the correlation metric more biologically meaningful, we subtract the mean unperturbed expression (µ^*ctrl*^) from the averaged predicted and observed gene expression profiles prior to computing the correlation-based metric. When applying Pearson correlation to these mean-centered profiles, we refer to this metric as Pearson delta (Pearson_Δ_), already used by previous studies^5,9,18^.

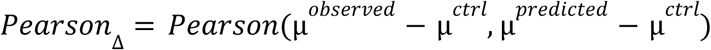

#### LogFC correlation

An alternative approach applies correlation-based metrics to log fold-change (logFC) profiles^26^. In this case, LogFC values are calculated between predicted perturbed and observed unperturbed cells, as well as between observed perturbed and unperturbed cells, and correlations are computed between the two resulting logFC profiles. Here, we employed Seurat’s^27^ *FindMarkers()* function to obtain the logFC values used in the logFC correlation metrics.

#### Maximum Mean Discrepancy

Maximum Mean Discrepancy (MMD) serves as a statistical measure to quantify distances between probability distributions. The (squared) MMD between two different distributions, *p* and *q*, is defined as:

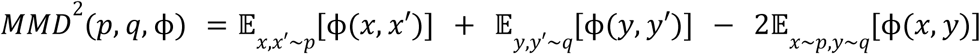

where ϕ is a kernel function, and *x* and *y* represent random variables from probability distributions *p* and *q*, respectively. A common choice for ϕ is the Gaussian Radial Basis Function (RBF) kernel, which introduces a bandwidth parameter (γ) to control sensitivity.

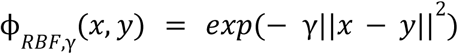

We considered γ equal to the inverse of the number of genes filtered in the final preprocessed datasets (i.e., γ = 1/5000). In practice, the expression profiles of single cells are considered as empirical distributions and the expectations in the MMD formula are calculated by averaging over pairs of points from each distribution.

#### Local MMD

In addition to the standard MMD formulation, we compute a modified metric referred to as Local MMD. This approach restricts the computation of pairwise distances to local neighborhoods rather than considering all possible pairs. By limiting comparisons to *k*-nearest neighbors in the feature space, Local MMD increases sensitivity to subtle, localized differences between distributions. This localized formulation enables more nuanced detection of distributional shifts that may be obscured when using the standard, fully global MMD metric. In contrast, when all pairwise distances are aggregated in the per-sample MMD calculation, the distinction between average in-distribution distances (within the same distribution) and out-distribution distances (from the opposite distribution) can be diluted.

For each point *x*_*i*_∼ *p* and *y*_*i*_∼ *q*, kernel values are calculated only with respect to their *k*-nearest neighbors from both their own distribution and the opposite distribution. Formally, (squared) Local MMD is defined as:

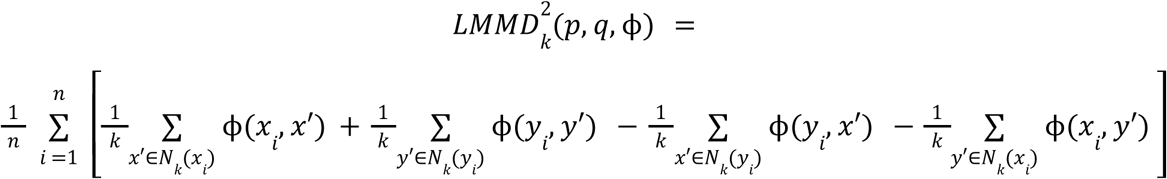

#### Local E-distance

When the kernel function in the MMD formulation is chosen as the negative Euclidean distance, the resulting metric is equivalent to the energy distance (E-distance). This kernel is defined as:

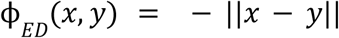

Unlike Gaussian kernels, the negative-distance kernel does not require a bandwidth parameter. We incorporated locality into the calculation of E-distance following a similar approach as described above, yielding the *Local E-distance*. Because the empirical computation of *MMD*^2^(*p*,*q*,ϕ)can yield negative values, we used the squared variants of MMD and E-distance (and their corresponding local variants) throughout the paper.

#### Wasserstein distance

The Wasserstein distance is a measure of distance between two probability distributions inspired from the optimal transport problem. Specifically, it calculates the minimum cost of transporting the probability mass of distribution µ such that the distribution ν is reconstructed. Formally, the p-Wasserstein is defined as:

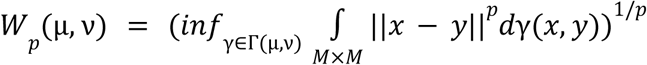

where Γ(µ, ν) is the set of all possible transport plans or joint distributions of µ and ν. Distributions µ and ν are defined on space *M*. γ is a joint probability on *M* × *M* whose marginals are µ and ν ^28^.

We used the Scipy’s *wasserstein_distance_nd()* function, which computes the Wasserstein-1 distance (*p* = 1) between two N-dimensional discrete distributions and uses the Euclidean norm as the cost function.

#### Mixing Index

We introduced the **Mixing Index**, a clustering-based metric that quantitatively evaluates the degree of co-distribution between predicted and true perturbed cells. The extent of integration between predicted and observed perturbed cells was evaluated in the presence of the ‘*whole dataset*’ comprising all cells subjected to the same perturbation across different cell types, together with unperturbed cells. This design ensures that predicted cells are evaluated for their alignment specifically with the true perturbed population, rather than with unrelated cell types or conditions.

Briefly, raw gene expression matrices were first log-normalized, centered and scaled prior to dimensionality reduction and clustering. Gene-wise scaling was performed using the mean and standard deviation computed across the *whole dataset*. Principal Component Analysis (PCA) was first fitted on the scaled *whole dataset* to capture the major sources of variation. The predicted and observed expression profiles of the target cell type were then projected into this precomputed PCA space to ensure a consistent embedding for mixing analysis.

Finally, the ‘*combined dataset*’ was clustered. The *combined dataset* included the predicted and observed profiles of the target perturbed cells in addition to the *whole dataset*. Clustering was performed using Seurat’s graph-based Louvain algorithm, implemented via the *FindNeighbors()* and *FindClusters()* functions (resolution = 2). Following clustering, the composition of each cluster was analyzed by counting the number of predicted cells and true observed cells corresponding to the target perturbation assigned to this cluster. Specifically, for each cluster *C_k_*, we defined 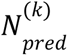 and 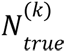 as the number of predicted and observed cells, respectively, and 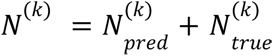 as the total number of cells in the given cluster. Note that each cluster may also contain cells other than the predicted and observed cells with the target perturbation, but their presence in the cluster has no impact on the mixing index computation. If the total number of predicted and observed cells of the target perturbation was equal, the mixing index for each cluster was then calculated as:

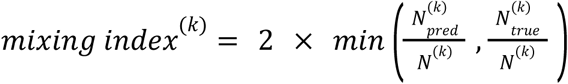

This formulation yields a value between 0 and 1, where 0 indicates complete segregation and 1 corresponds to perfect mixing (i.e., a 1:1 ratio of predicted and true cells). The overall mixing index was obtained by weighted averaging the per-cluster scores across all clusters.

### Noise sensitivity analysis

#### Gene-level correlation disruption (shuffling noise)

In each repeat, the true perturbed cells of a cell type were randomly divided into two equal subsets (Figure 6A): one designated as the ‘*test*’ split and the other as the ‘*noisy*’ split. To introduce controlled noise, we randomly selected a predefined percentage of genes (1% to 100%) and permuted their expression values across the cells within the noisy split. This procedure preserved per-gene marginal distributions but disrupted potential gene-gene dependencies and co-expression patterns within the noisy split. We then computed the divergence between each noisy split and the unchanged test split using distribution-based evaluation metrics such as mixing index, Wasserstein distance and E-distance. This process was repeated across several random splits to account for sampling variability.

The noisy splits could be constructed in either a *nested* or a *non-nested* manner. In the nested setting, higher noise levels were generated cumulatively: to obtain noise level *p*+*q*, the noisy split at level *p* was further shuffled for an additional *q*% of genes..

#### Expression-matrix noise addition

In a complementary strategy, we introduced randomized noise at the expression matrix level. In each repeat, the true perturbed single cells of a given cell type were split into a ‘*test*’ set and a ‘*noisy*’ set. We randomly selected a percentage of individual gene expression values across the entire matrix of the noisy split ranging from 1% to 100%. These selected values were then replaced with values sampled from the same gene but from a randomly selected single cell within a pool consisting of the mixture of other cell types under the same perturbation. This procedure introduces external noise into the gene expression structure of the noisy split. We then computed the divergence between the noisy and test splits using distribution-based evaluation metrics.

#### Sensitivity to noise

To quantify the sensitivity of a distribution-based metric to shuffling-noise, we defined a *sensitivity* criterion. Specifically, for a nested shuffling-noise experiment performed for several repeats, we computed

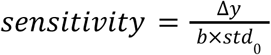

where *Δy* denotes the mean metric change from noise level 0 to 100, *std_0_* is the standard deviation at noise level 0 across sampling repeats and *b* is a constant that adjusts the magnitude of the sensitivity. In this study, we set *b* to 4.

### Generating synthetic datasets

To systematically evaluate the behavior of distribution-based metrics under controlled settings, we generated two classes of synthetic datasets.

#### Negative binomial RNA-seq simulation

We employed a custom single-cell RNA-seq simulator to generate count-based gene expression data that partially mimic the real transcriptomic profiles (described in detail in Supplementary Note 1). Briefly, cell populations were generated with expression counts for each gene sampled from a negative binomial distribution parameterized by user-specified distribution parameters. These parameters could alternatively be inferred from a real single-cell dataset, allowing the simulator to approximate gene-specific expression variance observed in real single-cell profiles.

To model different cell types or controlled perturbation effects, a defined fraction of genes was randomly selected and their expression levels were up- or down-regulated by specified fold-change factors. To imitate the high sparsity characteristic of real single-cell RNA-seq data, we additionally incorporated optional dropout mechanisms, in which a specified percentage of expression values was randomly masked to zero under various dropout scenarios originally described by Jiang et al.^29^ (Supplementary Note 2).

#### Multivariate normal distribution simulation

In addition, we generated synthetic datasets from a multivariate normal distribution. Each sample consisted of 5,000 features (representing genes), with desired number of data points (representing single cells).

In order to investigate the behavior of distribution-based metrics, initially two identically distributed samples were generated. To simulate increasing divergence between the two distributions, we progressively reduced the per-gene variance of one of the samples by scaling it with a factor less than 1 (from 0.9 to 0.1). This operation maintained the mean structure while reducing spread, leading to a more centralized cluster. We then evaluated how the Wasserstein distance, E-distance, and mixing index between the two samples varied in response to this change in distribution.

### Gene categories

To examine how gene selection influences evaluation outcomes, we categorized genes into three groups based on statistical significance of differential expression and the prevalence of zero expression:

**Trivial**: genes that are differentially expressed (p-value ≤ 0.05) and exhibit zero expression in all control cells (for upregulated DEGs) or in all perturbed cells (for downregulated DEGs). Genes were also labeled as *trivial* when the number of expressing cells differed by more than a tenfold factor between the two conditions.

**Non-trivial**: differentially expressed genes that do not meet the above criteria.

**Non-significant**: genes showing no statistically significant difference between perturbed and control conditions (p-value > 0.05)

### Benchmark and baseline models

#### Compositional Perturbation Autoencoder (CPA)

Combines interpretable linear modeling with adversarial training to learn disentangled latent representations of perturbations, covariates, and basal states. CPA encodes gene expression profiles into latent embeddings, removes perturbation and covariate effects, and then recombines these factors with the basal state to reconstruct perturbed profiles^21^.

#### scPRAM

Integrates attention mechanisms with optimal transport within a Variational Autoencoder (VAE) architecture to predict perturbation responses. Training data are encoded into a latent space, and unpaired cells are aligned before and after perturbation via optimal transport (Sinkhorn algorithm) to estimate cell-wise perturbation vectors.

For a test cell, scPRAM uses an attention mechanism to infer its perturbation response: the latent representation of the test cell serves as a query, while the latent embeddings of pre-perturbed cells in the training set act as keys and their corresponding optimal-transport-derived perturbation vectors serve as values. The attention-weighted aggregation of these vectors yields a predicted perturbation shift, which is then added to the test cell’s latent embedding and decoded to reconstruct the perturbed profiles ^7^.

#### CAE

We implemented a Conditional Autoencoder (CAE), a type of autoencoder that incorporates cell conditions (e.g., cell type and perturbation) as one-hot encoded vectors concatenated with both input gene expression and the latent space vector^30,31^. The CAE architecture included hidden layers of 512 and 256 units, a latent dimension of 50, a dropout rate of 0.2, and ReLU activation.

#### No-perturb

Assumes that perturbations have no effect, predicting post-perturbation gene expression profiles identical to the unperturbed profiles.

#### LogFC-transfer

Computes fold change profiles between perturbed and control conditions in each non-target cell type using Seurat’s *FIndMarkers()*, then multiplies the average calculated logFC by the control gene expression profiles of the target cell type to predict the perturbed profiles.

#### Random-perturb

Generates perturbed single cells by randomly sampling cells from other cell types subjected to the same perturbation.

## Data availability

All datasets used in this study are publicly available. For the Kang^23^ dataset (GSE96583), we used the preprocessed data provided by Lotfollahi *et al*.^21^ at [Google Drive](https://drive.google.com/drive/u/0/folders/1yFB0gBr72_KLLp1asojxTgTqgz6cwpju). The count data corresponding to the Datlinger^24^ dataset was obtained from GEO accession GSM5151370. The Sciplex3^25^ dataset (GSM4150378) was available via scPerturb^3^ at https://doi.org/10.5281/zenodo.13350497 (ref.^32^).

## Code availability

The source code corresponding to the CrossSplit pipeline is available on GitHub at https://github.com/scPerturbationStudies/scPerturb-eval

## Acknowledgements

This work is based upon research funded by Iran National Science Foundation (INSF) under project No. 4032273. The funder had no role in study design, data collection and analysis, decision to publish, or preparation of the manuscript.

## Contributions

M.K., M.H., H.M., and S.S. conceived the project. M.H., M.K., and H.M. developed the methodology. M.H. and M.K. implemented the pipeline and methods in Python and conducted all experiments and visualizations. M.K. collected and preprocessed the data. H.M. supervised the study and verified the findings. M.H. and M.K. wrote the first draft of the manuscript with feedback from H.M. M.H. finalized the last version of manuscript and figures. All authors reviewed and approved the final version of the manuscript.

## Competing interests

The authors declare no competing interests.

## Notes

### Competing Interest Statement

The authors have declared no competing interest.

## References

1. Heumos, L. et al. Best practices for single-cell analysis across modalities. Nat Rev Genet 24, 550–572 (2023).

2. Jovic, D. et al. Single-cell RNA sequencing technologies and applications: A brief overview. Clin Transl Med 12, e694 (2022).

3. Peidli, S. et al. scPerturb: harmonized single-cell perturbation data. Nat Methods 21, 531–540 (2024).

4. Lotfollahi, M., Wolf, F. A. & Theis, F. J. scGen predicts single-cell perturbation responses. Nat Methods 16, 715–721 (2019).

5. Roohani, Y., Huang, K. & Leskovec, J. Predicting transcriptional outcomes of novel multigene perturbations with GEARS. Nat Biotechnol 42, 927–935 (2024).

6. Bunne, C. et al. Learning single-cell perturbation responses using neural optimal transport. Nat Methods 20, 1759–1768 (2023).

7. Jiang, Q., Chen, S., Chen, X. & Jiang, R. scPRAM accurately predicts single-cell gene expression perturbation response based on attention mechanism. Bioinformatics 40, (2024).

8. Hao, M. et al. Large-scale foundation model on single-cell transcriptomics. Nat Methods 21, 1481–1491 (2024).

9. Cui, H. et al. scGPT: toward building a foundation model for single-cell multi-omics using generative AI. Nat Methods 21, 1470–1480 (2024).

10. Li, L. et al. A Systematic Comparison of Single-Cell Perturbation Response Prediction Models. bioRxiv 2024.12.23.630036 (2024) doi:10.1101/2024.12.23.630036.

11. Ahlmann-Eltze, C., Huber, W. & Anders, S. Deep-learning-based gene perturbation effect prediction does not yet outperform simple linear baselines. Nature Methods 22, 1657–1661 (2025).

12. Yang, F. et al. scBERT as a large-scale pretrained deep language model for cell type annotation of single-cell RNA-seq data. Nature Machine Intelligence 4, 852–866 (2022).

13. Theodoris, C. V. et al. Transfer learning enables predictions in network biology. Nature 618, 616–624 (2023).

14. Wenteler, A., et al. PertEval-scFM: Benchmarking Single-Cell Foundation Models for Perturbation Effect Prediction. bioRxiv 2024.10.02.616248 (2024) doi:10.1101/2024.10.02.616248.

15. Wu, Y., et al. PerturBench: Benchmarking Machine Learning Models for Cellular Perturbation Analysis. (2024).

16. Csendes, G., Sanz, G., Szalay, K. Z. & Szalai, B. Benchmarking foundation cell models for post-perturbation RNA-seq prediction. BMC Genomics 26, 393 (2025).

17. Kedzierska, K. Z., Crawford, L., Amini, A. P. & Lu, A. X. Zero-shot evaluation reveals limitations of single-cell foundation models. Genome Biology 26, 101 (2025).

18. Viñas Torné, R., et al. Systema: a framework for evaluating genetic perturbation response prediction beyond systematic variation. Nature Biotechnology 1–10 (2025).

19. Miller, H. E. et al. Deep learning-based genetic perturbation models *do* outperform uninformative baselines on well-calibrated metrics. bioRxiv 2025.10.20.683304 (2025) doi:10.1101/2025.10.20.683304.

20. Mejia, G. M. et al. Diversity by design: Addressing mode collapse improves scRNA-seq perturbation modeling on well-calibrated metrics. arXiv [q-bio.GN] (2025).

21. Lotfollahi, M. et al. Predicting cellular responses to complex perturbations in high-throughput screens. Molecular Systems Biology (2023) doi:10.15252/msb.202211517.

22. Rich, J. M., et al. The impact of package selection and versioning on single-cell RNA-seq analysis. bioRxivorg (2024) doi:10.1101/2024.04.04.588111.

23. Kang, H. M. et al. Multiplexed droplet single-cell RNA-sequencing using natural genetic variation. Nat Biotechnol 36, 89–94 (2018).

24. Datlinger, P. et al. Ultra-high-throughput single-cell RNA sequencing and perturbation screening with combinatorial fluidic indexing. Nat Methods 18, 635–642 (2021).

25. Srivatsan, S. R. et al. Massively multiplex chemical transcriptomics at single-cell resolution. Science 367, 45–51 (2020).

26. Qi, X. et al. Predicting transcriptional responses to novel chemical perturbations using deep generative model for drug discovery. Nat Commun 15, 9256 (2024).

27. Satija, R., Farrell, J. A., Gennert, D., Schier, A. F. & Regev, A. Spatial reconstruction of single-cell gene expression data. Nat. Biotechnol. 33, 495–502 (2015).

28. Panaretos, V. M. & Zemel, Y. Statistical aspects of Wasserstein distances. Annu. Rev. Stat. Appl. 6, 405–431 (2019).

29. Jiang, R., Sun, T., Song, D. & Li, J. J. Statistics or biology: the zero-inflation controversy about scRNA-seq data. Genome Biol 23, 31 (2022).

30. Hinton, G. E. & Salakhutdinov, R. R. Reducing the dimensionality of data with neural networks. Science 313, 504–507 (2006).

31. Sohn, K., Lee, H. & Yan, X. Learning structured output representation using deep conditional generative models. Neural Inf Process Syst 28, 3483–3491 (2015).

32. Peidli, S., et al. scPerturb Single-Cell Perturbation Data: RNA and protein h5ad files. Zenodo 10.5281/ZENODO.13350497 (2022).

