## Supplementary Information for "Evaluating Single-Cell Perturbation Response Models Is Far from Straightforward"

---

[1] Department of Bioinformatics, Institute of Biochemistry and Biophysics, University of Tehran, Tehran, Iran
[2] Independent scholar

[*] Equal contribution

#### Supplementary Note 1- Simulation model

We developed a simulation framework designed to generate scRNA-seq count data with *N* cells and *M* genes under different experimental conditions.We begin by defining the key experimental variables: $K$, the number of conditions influencing gene expression profiles, (e.g., $K$ could represent variables like *cell type*, *dosage*, or *perturbation*) and $L_k$, the number of levels for each condition $k$ (e.g., for the condition *cell type*, $L_k$ could represent the levels 'B cells', 'T cells', 'Monocytes', etc.). For a single cell $i$, the condition is denoted by a vector

$$c_i = (c_i^1, . . . , c_i^K)$$

where $c_i^k$ represents the level of condition $k$ for cell $i$. The expression vector for a single cell $i$ is represented as

$$E_i = (E_i^1, . . . , E_i^M).$$

We assume that the baseline expression $E_{i,b}^m$ for gene $m$ is sampled from a negative binomial distribution,

$$E^m_{i,b} \sim NB(r^m, p^m)$$

where $r^m$ and $p^m$ are the parameters of negative binomial distribution for gene $m$. The parameter $p^m$ is defined as a function of the baseline mean of the distribution, $\lambda^m_b$, and $r^m$,

$$p^m = \frac{r^m}{\lambda^m_b + r^m}$$

The baseline mean expression $\lambda^m_b$ is modeled to capture inherent biological variability across genes. It can be simulated from a uniform distribution, e.g.,:

$$\lambda^m_b \sim Uniform(1, 10)$$

or, can be adjusted to account for gene length, from the following expression:

$$\lambda^m_b = \frac{Length_m}{\sum_{j=1}^{M} Length_j} \times counts_{total}$$

where $counts_{total}$ is a desired value for the total read counts in a single cell. The baseline mean expression can also be estimated from a real expression dataset. We model condition-specific effects using a modulation factor $B^m_{c^k_i}$, where $B^m_{c^k_i} = 1$ indicates no effect, $B^m_{c^k_i} > 1$ up-regulation and $B^m_{c^k_i} < 1$ down regulation. Gene expression under condition vector $c_i$ is defined as follows

$$E^m_i = E^m_{i,b} \prod_{k=1}^{K} B^m_{c^k_i}$$

For each condition $k$, a fraction $f^k$ of genes is affected. Modulation factors for affected genes are sampled with equal probability from $B^m_{c^k_i} \sim Uniform(2, 5)$ for up-regulation or $B^m_{c^k_i} \sim Uniform(\frac{1}{5}, \frac{1}{2})$ for down-regulation.

In the final step, to simulate technical dropout effects commonly observed in scRNA-seq data, we apply a dropout model to the simulated expression matrix by randomly setting a proportion of non-zero entries to zero, following the masking schemes described by

Jiang *et al.*[1] Dropout probabilities are either estimated from real data or fixed to a constant value.

#### Supplementary Note 2- Masking Scenarios

Detailed definitions of the masking schemes could be found in the original study[1]. Let $p$ denote the overall dropout probability and $p_i$ the gene-specific dropout probability for gene $i$. Let $n$ be the total number of non-zero expression values across all genes, and $n_i$ the number of non-zero expression values for gene $i$. Here, we briefly summarize each scenario:

**random-all**: randomly masks $N \sim Binomial(n, p)$ non-zero counts across the entire expression matrix.

**quantile-all**: masks the smallest $[np]$ non-zero counts across the entire expression matrix.

**quantile-same**: for each gene $i$, masks the smallest $\left[n_i p\right]$ non-zero counts of that gene.

**random-gene-specific**: for each gene $i$, randomly masks $N_i \sim Binomial(n_i, p_i)$ non-zero counts. The gene-specific dropout probability $p_i$ is defined to decrease exponentially with the mean non-zero expression level of gene $i$ or $\mu_i$ (i.e., $p_i = exp(-\lambda\mu_i^2)$).

**quantile-gene-specific**: for each gene $i$, masks the smallest $\left[n_i p_i\right]$ non-zero counts, where $p_i$ is defined in the same manner as in the random-gene-specific scheme.

#### Supplementary Note 3- Evaluating DEG Recovery Methods

We evaluated multiple statistical methods for detecting differentially expressed genes (DEGs) in perturbation experiments. For this, we leveraged our simulator to generate perturbed and unperturbed single-cell data under controlled dropout scenarios, as defined in Jiang *et al.*[1], with desired dropout probabilities. The condition-specific modulation factors $B^{m_k}_{c_i}$ are known during simulation, allowing us to define ground-truth.

We compared four DEG-detection approaches that compute p-values for differential expression between perturbed and unperturbed cells:

**Wilcoxon test (with zeros)**: the standard Wilcoxon rank-sum test applied directly to the simulated log-normalized data, using all expression values - including zeros. For each gene, we used Scipy's *ranksums()* function, applying the '*less*' and '*greater*' alternatives to detect down- and up-regulated genes, respectively.

**Wilcoxon test (non-zero)**: the Wilcoxon rank-sum test applied after removing cells with zero expression for the gene under consideration. If a gene had zero expression across all cells in either condition (perturbed or unperturbed), we applied a Wilcoxon signed-rank test instead.

**Seurat-MAST**: a method specifically designed to account for zero inflation in single-cell data[2], implemented in Seurat. We used Seurat's *FindMarkers()* function with *test.use="MAST"*.

**Seurat-Wilcoxon**: Seurat's default Wilcoxon implementation used in Seurat's *FindMarkers()*.

We simulated single-cell perturbation datasets across different dropout probabilities and dropout scenarios. The maximum dropout probability was selected such that the resulting proportion of zero expression values in the dataset matched that of the real Kang dataset ($\approx 95\%$). For each method, genes were ranked by statistical significance (perturbed vs. unperturbed), and recovery of true DEGs was quantified among the top-ranked genes.

Our analysis showed that increasing dropout probability led to reduced DEG-recovery performance overall, although the magnitude of decline varied by the method and the masking scheme (Supplementary Figure 10). Notably, Seurat-MAST consistently outperformed the other approaches across different dropout scenarios, demonstrating superior robustness to high sparsity. Based on these results, we selected Seurat-MAST as the primary DEG identification method for all downstream analyses.

#### Supplementary Note 4- PID evaluations across varying sample sizes

As detailed in the *Evaluation framework* section of the *Methods*, evaluation at each PID level was performed using a fixed *comparison size* equal to half the number of true target perturbed cells. Generative models were required to produce the same number of predictions to remove the influence of varying sample sizes across PID levels. However, this procedure necessitates up-sampling for the reference group at lower PID percentages, potentially introducing repeated cells (since the explored PID percentages were at most 50%, the size of the reference group never exceeded the comparison size).

To avoid introducing repeated cells in the reference group and at the same time, to explicitly assess the impact of sample size on metric values, we repeated the analysis using variable comparison sizes limited to the number of *seen* target perturbed cells at each PID percentage. An example of this analysis for B cells in the Kang dataset, using E-distance and mixing index, is shown in Supplementary Figure 5.

Comparing evaluation scores of each model across varying sample sizes at fixed PID levels revealed that E-distance remained largely stable, aside from minor fluctuations. In contrast, the mixing index exhibited sensitivity to sample size in certain cases; for example, the CAE model showed a pronounced decline in mixing index as the comparison size increased at a fixed PID level. These observations indicate that the mixing index can be influenced by sample size, a factor that should cautiously be taken into account when interpreting the results. Across all PID percentages and sample sizes, CPA consistently underperformed baseline models in terms of E-distance and yielded near-zero mixing index values. Similarly, scPRAM performed comparably to the no-perturb baseline on E-distance but consistently underperformed it on the mixing index.

#### Supplementary Note 5- Cross-group mixing experiment (nearest-neighbor analysis)

The Wasserstein distance between two probability distributions is often interpreted as the minimal “work” required to transform one distribution into the other. The overall cost is defined as the product of the amount of probability mass transported and the distance over which it is moved. Because the Euclidean norm is typically used as the ground distance in Wasserstein calculations, examining nearest-neighbor relationships between samples can provide insight into the counterintuitive behavior of Wasserstein observed under variance scaling.

We simulated two high-dimensional samples (5,000 genes, 100 cells each), drawn either from a multivariate normal distribution with zero mean or from a negative binomial distribution (mean = 100), with negative binomial counts subsequently log-normalized to a total sum of $10^4$. In the multivariate normal setting, sample 1 had identity covariance and in the negative binomial setting, the number of successes for sample 1 was $r = 1$. In both simulation schemes, sample 1 had fixed per-gene variance (scale = 1.0), whereas the variance of sample 2 was progressively reduced from 1.0 down to 0.91. For each configuration, we quantified the fraction of cells whose nearest neighbor belonged to the other sample.

As shown in Figures 5E and 5I, when sample 2 was slightly more compact (variance = 0.91 relative to sample 1), all of its cells' nearest neighbors were within its own group (2→1 (1-NN, %) = 0), while all cells from sample 1 found their nearest neighbors in sample 2 (1→2 (1-NN, %) = 100). This asymmetry gradually disappeared as the variance of sample 2 increased toward that of sample 1 (scaling coefficient → 1), converging to approximately equal nearest-neighbor counts in both directions. These asymmetric nearest-neighbor relationships help explain why Wasserstein distance can decrease as one distribution becomes overly concentrated in high-dimensional settings.

#### Supplementary Figures

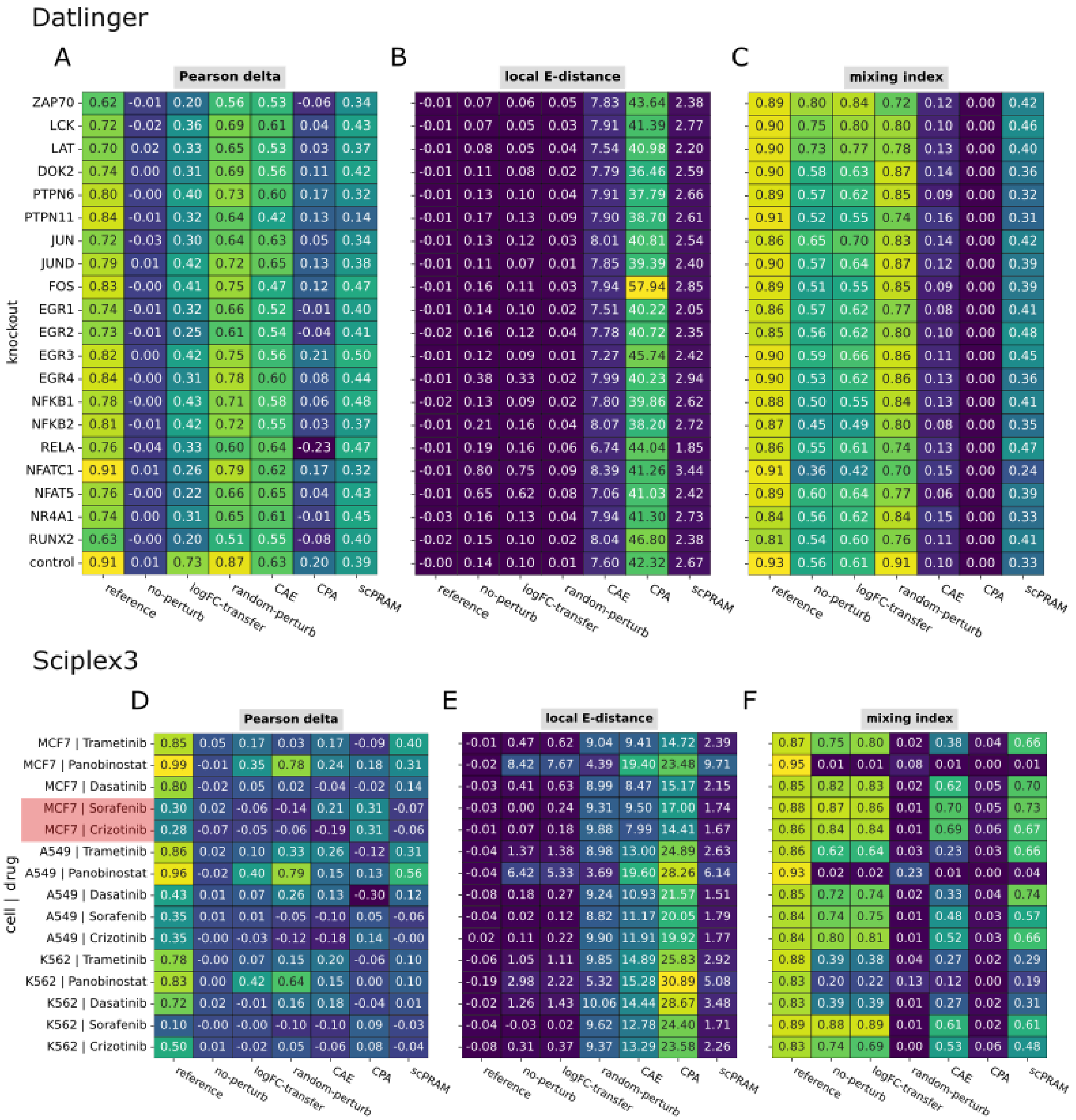


**Supplementary Figure 1. CrossSplit OOD evaluation results.** Heatmaps summarizing CrossSplit OOD evaluation results on Pearson delta, local E-distance, and mixing index across **(A-C)** the stimulated state of all gene knockouts of the Jurkat cell line in the Datlinger dataset and **(D-F)** all three cell lines in the Sciplex3 dataset treated with a selection of five drugs. Each model was trained to predict the perturbed state of each cell type under the Out-of-Distribution (OOD) setting. Values represent average performance across CrossSplit repetitions for baseline and deep learning models, as well as the reference model. Cases in which the reference model shows inferior performance relative to other models are highlighted in red.

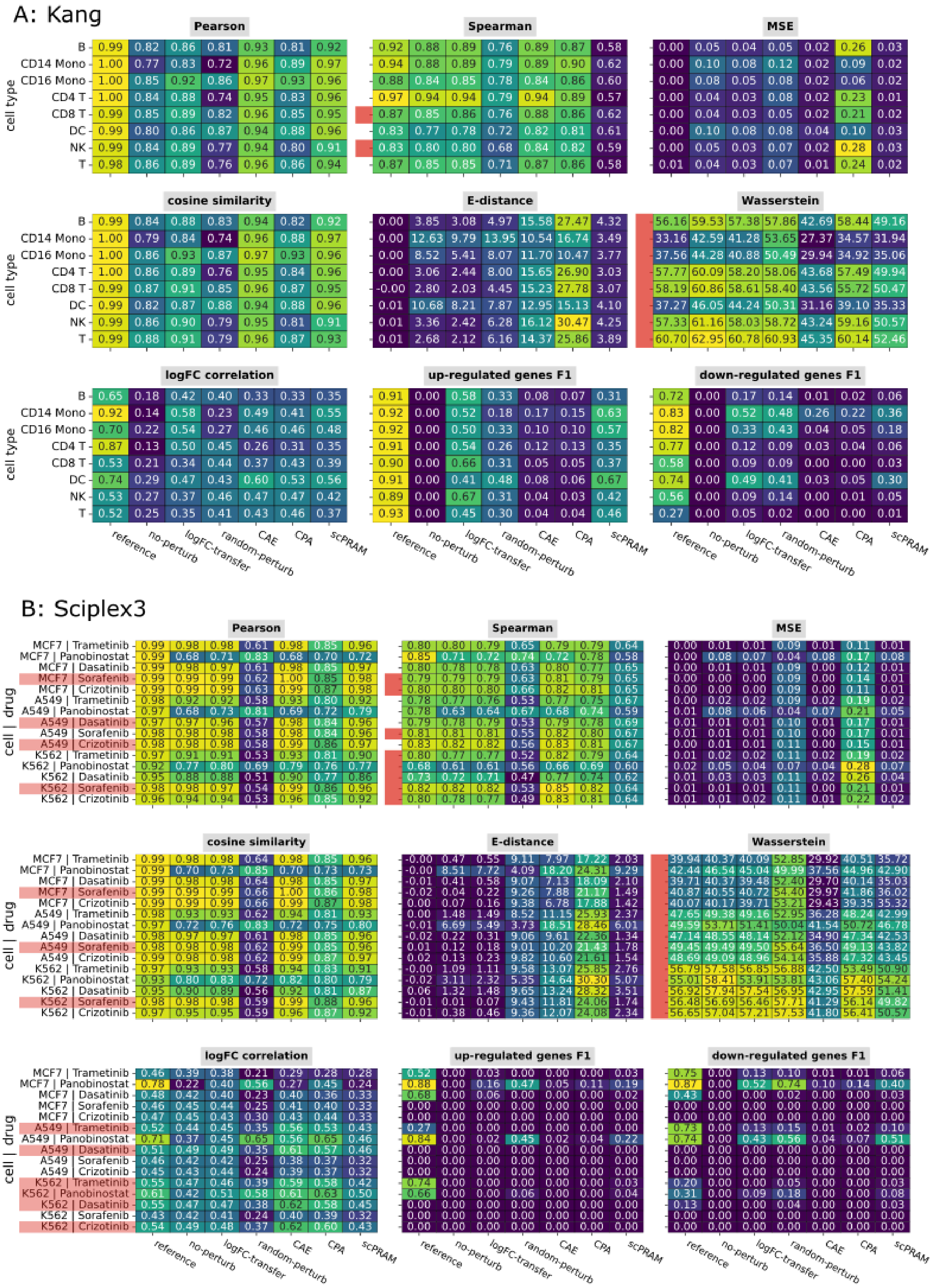
A: Kang
Pearson
Spearman
MSE
cosine similarity
E-distance
Wasserstein
logFC correlation
up-regulated genes F1
down-regulated genes F1
cell type
B
CD14 Mono
CD16 Mono
CD4 T
CD8 T
DC
NK
T
reference
no-perturb
logFC-transfer
random-perturb
CAE
CPA
scPRAM
B: Sciplex3
cell | drug
MCF7 | Trametinib
MCF7 | Panobinostat
MCF7 | Dasatinib
MCF7 | Sorafenib
MCF7 | Crizotinib
A549 | Trametinib
A549 | Panobinostat
A549 | Dasatinib
A549 | Sorafenib
A549 | Crizotinib
K562 | Trametinib
K562 | Panobinostat
K562 | Dasatinib
K562 | Sorafenib
K562 | Crizotinib

**Supplementary Figure 2. CrossSplit OOD evaluation results.** Heatmaps summarizing CrossSplit OOD evaluation results based on several common but less reliable metrics across **(A)** the stimulated state of all cell types in the Kang dataset and **(B)** all three cell lines in the Sciplex3 dataset treated with a selection of five drugs. Each model was trained to predict the stimulated state of each cell type under the Out-of-Distribution (OOD) setting. Values represent the average performance across CrossSplit repetitions for baseline, deep learning and the reference models. Cases where the reference model shows inferior performance relative to other models are highlighted in red.

### Datlinger

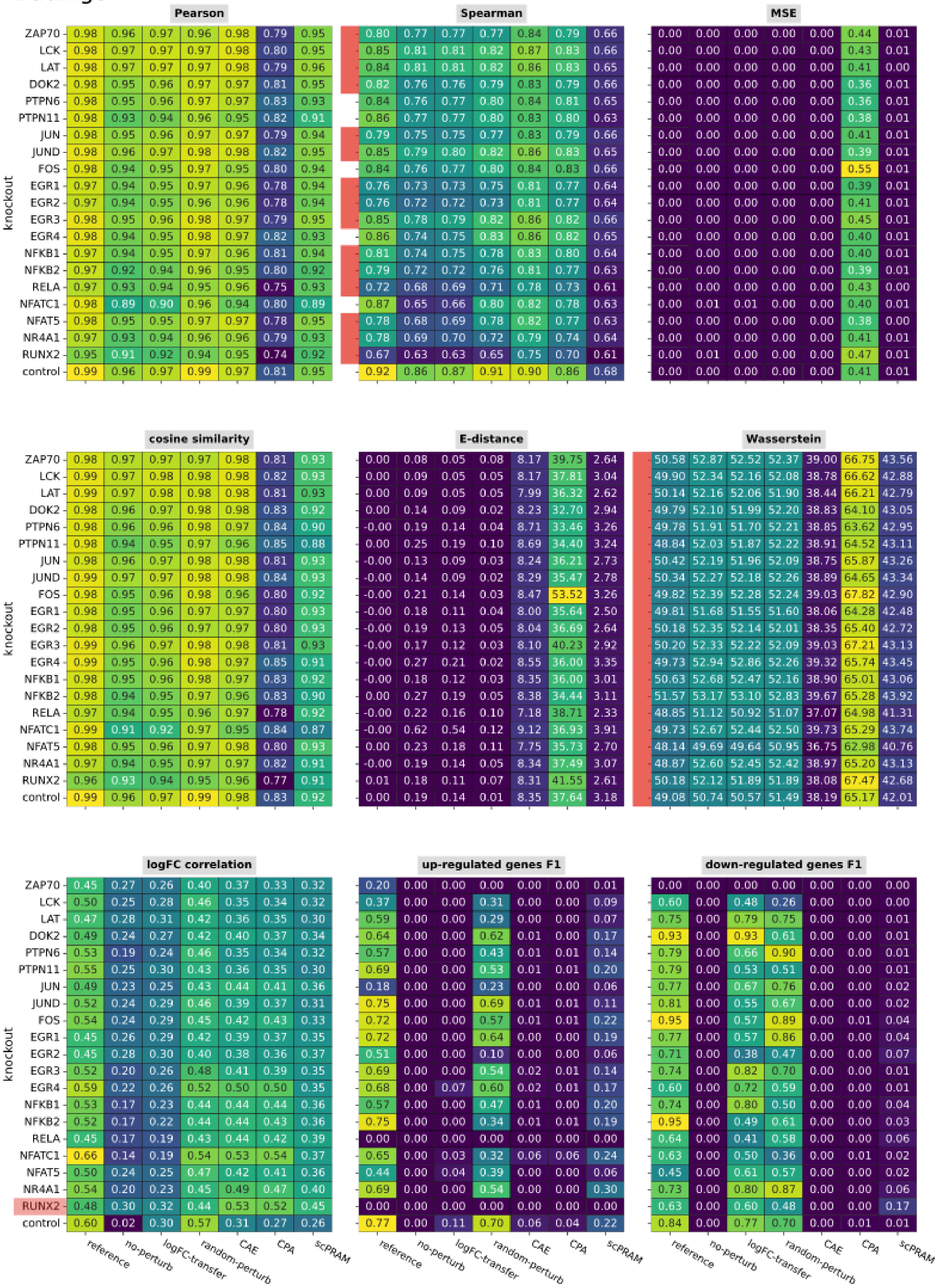
Pearson
ZAP70 0.98 0.96 0.97 0.96 0.98 0.79 0.95
LCK 0.98 0.97 0.97 0.97 0.98 0.80 0.95
LAT 0.98 0.97 0.97 0.97 0.98 0.79 0.96
DOK2 0.98 0.95 0.96 0.97 0.97 0.81 0.95
PTPN6 0.98 0.95 0.96 0.97 0.97 0.83 0.93
PTPN11 0.98 0.93 0.94 0.96 0.95 0.82 0.91
JUN 0.98 0.95 0.96 0.97 0.97 0.79 0.94
JUND 0.98 0.96 0.97 0.98 0.98 0.82 0.95
FOS 0.98 0.94 0.95 0.97 0.95 0.80 0.94
EGR1 0.97 0.94 0.95 0.97 0.96 0.78 0.94
EGR2 0.97 0.94 0.95 0.96 0.96 0.78 0.94
EGR3 0.98 0.95 0.96 0.98 0.97 0.79 0.95
EGR4 0.98 0.94 0.95 0.98 0.97 0.82 0.93
NFKB1 0.97 0.94 0.95 0.97 0.96 0.81 0.94
NFKB2 0.97 0.92 0.94 0.96 0.95 0.80 0.92
RELA 0.97 0.93 0.94 0.95 0.96 0.75 0.93
NFATC1 0.98 0.89 0.90 0.96 0.94 0.80 0.89
NFAT5 0.98 0.95 0.95 0.97 0.97 0.78 0.95
NR4A1 0.97 0.93 0.94 0.96 0.96 0.79 0.93
RUNX2 0.95 0.91 0.92 0.94 0.95 0.74 0.92
control 0.99 0.96 0.97 0.99 0.97 0.81 0.95
Spearman
0.80 0.77 0.77 0.77 0.84 0.79 0.66
0.85 0.81 0.81 0.82 0.87 0.83 0.66
0.84 0.81 0.81 0.82 0.86 0.83 0.65
0.82 0.76 0.76 0.79 0.83 0.79 0.66
0.84 0.76 0.77 0.80 0.84 0.81 0.65
0.86 0.77 0.77 0.80 0.83 0.80 0.63
0.79 0.75 0.75 0.77 0.83 0.79 0.66
0.85 0.79 0.80 0.82 0.86 0.83 0.65
0.84 0.76 0.77 0.80 0.84 0.83 0.66
0.76 0.73 0.73 0.75 0.81 0.77 0.64
0.76 0.72 0.72 0.73 0.81 0.77 0.64
0.85 0.78 0.79 0.82 0.86 0.82 0.66
0.86 0.74 0.75 0.83 0.86 0.82 0.65
0.81 0.74 0.75 0.78 0.83 0.80 0.64
0.79 0.72 0.72 0.76 0.81 0.77 0.63
0.72 0.68 0.69 0.71 0.78 0.73 0.61
0.87 0.65 0.66 0.80 0.82 0.78 0.63
0.78 0.68 0.69 0.78 0.82 0.77 0.63
0.78 0.69 0.70 0.72 0.79 0.74 0.64
0.67 0.63 0.63 0.65 0.75 0.70 0.61
0.92 0.86 0.87 0.91 0.90 0.86 0.68
MSE
0.00 0.00 0.00 0.00 0.00 0.44 0.01
0.00 0.00 0.00 0.00 0.00 0.43 0.01
0.00 0.00 0.00 0.00 0.00 0.41 0.00
0.00 0.00 0.00 0.00 0.00 0.36 0.01
0.00 0.00 0.00 0.00 0.00 0.36 0.01
0.00 0.00 0.00 0.00 0.00 0.38 0.01
0.00 0.00 0.00 0.00 0.00 0.41 0.01
0.00 0.00 0.00 0.00 0.00 0.39 0.01
0.00 0.00 0.00 0.00 0.00 0.55 0.01
0.00 0.00 0.00 0.00 0.00 0.39 0.01
0.00 0.00 0.00 0.00 0.00 0.41 0.01
0.00 0.00 0.00 0.00 0.00 0.45 0.01
0.00 0.00 0.00 0.00 0.00 0.40 0.01
0.00 0.00 0.00 0.00 0.00 0.40 0.01
0.00 0.00 0.00 0.00 0.00 0.39 0.01
0.00 0.00 0.00 0.00 0.00 0.43 0.00
0.00 0.01 0.01 0.00 0.00 0.40 0.01
0.00 0.00 0.00 0.00 0.00 0.38 0.00
0.00 0.00 0.00 0.00 0.00 0.41 0.01
0.00 0.01 0.00 0.00 0.00 0.47 0.01
0.00 0.00 0.00 0.00 0.00 0.41 0.01
knockout
cosine similarity
ZAP70 0.98 0.97 0.97 0.97 0.98 0.81 0.93
LCK 0.99 0.97 0.98 0.98 0.98 0.82 0.93
LAT 0.99 0.97 0.98 0.98 0.98 0.81 0.93
DOK2 0.98 0.96 0.97 0.98 0.98 0.83 0.92
PTPN6 0.98 0.96 0.96 0.98 0.97 0.84 0.90
PTPN11 0.98 0.94 0.95 0.97 0.96 0.85 0.88
JUN 0.98 0.96 0.97 0.98 0.98 0.81 0.93
JUND 0.99 0.97 0.97 0.98 0.98 0.84 0.93
FOS 0.98 0.95 0.96 0.98 0.96 0.80 0.92
EGR1 0.98 0.95 0.96 0.97 0.97 0.80 0.93
EGR2 0.98 0.95 0.96 0.97 0.97 0.80 0.93
EGR3 0.99 0.96 0.97 0.98 0.98 0.81 0.93
EGR4 0.99 0.95 0.96 0.98 0.97 0.85 0.91
NFKB1 0.98 0.95 0.96 0.98 0.97 0.83 0.92
NFKB2 0.98 0.94 0.95 0.97 0.96 0.83 0.90
RELA 0.97 0.94 0.95 0.96 0.97 0.78 0.92
NFATC1 0.99 0.91 0.92 0.97 0.95 0.84 0.87
NFAT5 0.98 0.95 0.96 0.97 0.98 0.80 0.93
NR4A1 0.97 0.94 0.95 0.97 0.97 0.82 0.91
RUNX2 0.96 0.93 0.94 0.95 0.96 0.77 0.91
control 0.99 0.96 0.97 0.99 0.98 0.83 0.92
E-distance
0.00 0.08 0.05 0.08 8.17 39.75 2.64
0.00 0.09 0.05 0.05 8.17 37.81 3.04
0.00 0.09 0.05 0.05 7.99 36.32 2.62
0.00 0.14 0.09 0.02 8.23 32.70 2.94
-0.00 0.19 0.14 0.04 8.71 33.46 3.26
0.00 0.25 0.19 0.10 8.69 34.40 3.24
-0.00 0.13 0.09 0.03 8.24 36.21 2.73
-0.00 0.14 0.09 0.02 8.29 35.47 2.78
-0.00 0.21 0.14 0.03 8.47 53.52 3.26
-0.00 0.18 0.11 0.04 8.00 35.64 2.50
-0.00 0.19 0.13 0.05 8.04 36.69 2.64
-0.00 0.17 0.12 0.03 8.10 40.23 2.92
-0.00 0.27 0.21 0.02 8.55 36.00 3.35
-0.00 0.18 0.12 0.03 8.35 36.00 3.01
0.00 0.27 0.19 0.05 8.38 34.44 3.11
-0.00 0.22 0.16 0.10 7.18 38.71 2.33
-0.00 0.62 0.54 0.12 9.12 36.93 3.91
0.00 0.23 0.18 0.11 7.75 35.73 2.70
-0.00 0.19 0.14 0.05 8.34 37.49 3.07
0.01 0.18 0.11 0.07 8.31 41.55 2.61
0.00 0.19 0.14 0.01 8.35 37.64 3.18
Wasserstein
50.58 52.87 52.52 52.37 39.00 66.75 43.56
49.90 52.34 52.16 52.08 38.78 66.62 42.88
50.14 52.16 52.06 51.90 38.44 66.21 42.79
49.79 52.10 51.99 52.20 38.83 64.10 43.05
49.78 51.91 51.70 52.21 38.85 63.62 42.95
48.84 52.03 51.87 52.22 38.91 64.52 43.11
50.42 52.19 51.96 52.09 38.75 65.87 43.26
50.34 52.27 52.18 52.26 38.89 64.65 43.34
49.82 52.39 52.28 52.24 39.03 67.82 42.90
49.81 51.68 51.55 51.60 38.06 64.28 42.48
50.18 52.35 52.14 52.01 38.35 65.40 42.72
50.20 52.33 52.22 52.09 39.03 67.21 43.13
49.73 52.94 52.86 52.26 39.32 65.74 43.45
50.63 52.68 52.47 52.16 38.90 65.01 43.06
51.57 53.17 53.10 52.83 39.67 65.28 43.92
48.85 51.12 50.92 51.07 37.07 64.98 41.31
49.73 52.67 52.44 52.50 39.73 65.29 43.74
48.14 49.69 49.64 50.95 36.75 62.98 40.76
48.87 52.60 52.45 52.42 38.97 65.20 43.13
50.18 52.12 51.89 51.89 38.08 67.47 42.68
49.08 50.74 50.57 51.49 38.19 65.17 42.01
knockout
logFC correlation
ZAP70 0.45 0.27 0.26 0.40 0.37 0.33 0.32
LCK 0.50 0.25 0.28 0.46 0.35 0.34 0.32
LAT 0.47 0.28 0.31 0.42 0.36 0.35 0.30
DOK2 0.49 0.24 0.27 0.42 0.40 0.37 0.34
PTPN6 0.53 0.19 0.24 0.46 0.35 0.34 0.32
PTPN11 0.55 0.25 0.30 0.43 0.36 0.35 0.30
JUN 0.49 0.23 0.25 0.43 0.44 0.41 0.36
JUND 0.52 0.24 0.29 0.46 0.39 0.37 0.31
FOS 0.54 0.24 0.29 0.45 0.42 0.43 0.33
EGR1 0.45 0.26 0.29 0.42 0.39 0.37 0.35
EGR2 0.45 0.28 0.30 0.40 0.38 0.36 0.37
EGR3 0.52 0.20 0.26 0.48 0.41 0.39 0.35
EGR4 0.59 0.22 0.26 0.52 0.50 0.50 0.35
NFKB1 0.53 0.17 0.23 0.44 0.44 0.44 0.36
NFKB2 0.52 0.17 0.22 0.44 0.44 0.43 0.36
RELA 0.45 0.17 0.19 0.43 0.44 0.42 0.39
NFATC1 0.66 0.14 0.19 0.54 0.53 0.54 0.37
NFAT5 0.50 0.24 0.25 0.47 0.42 0.41 0.36
NR4A1 0.54 0.20 0.23 0.45 0.49 0.47 0.40
RUNX2 0.48 0.30 0.32 0.44 0.53 0.52 0.45
control 0.60 0.02 0.30 0.57 0.31 0.27 0.26
up-regulated genes F1
0.20 0.00 0.00 0.00 0.00 0.00 0.01
0.37 0.00 0.00 0.31 0.00 0.00 0.09
0.59 0.00 0.00 0.29 0.00 0.00 0.07
0.64 0.00 0.00 0.62 0.01 0.00 0.17
0.57 0.00 0.00 0.43 0.01 0.01 0.14
0.69 0.00 0.00 0.53 0.01 0.01 0.20
0.18 0.00 0.00 0.23 0.00 0.00 0.06
0.75 0.00 0.00 0.69 0.01 0.01 0.11
0.72 0.00 0.00 0.57 0.01 0.01 0.22
0.72 0.00 0.00 0.64 0.00 0.00 0.19
0.51 0.00 0.00 0.10 0.00 0.00 0.06
0.69 0.00 0.00 0.54 0.02 0.01 0.14
0.68 0.00 0.07 0.60 0.02 0.01 0.17
0.57 0.00 0.00 0.47 0.01 0.00 0.20
0.75 0.00 0.00 0.34 0.01 0.01 0.19
0.00 0.00 0.00 0.00 0.00 0.00 0.00
0.65 0.00 0.03 0.32 0.06 0.06 0.24
0.44 0.00 0.04 0.39 0.00 0.00 0.06
0.69 0.00 0.00 0.54 0.00 0.00 0.30
0.00 0.00 0.00 0.00 0.00 0.00 0.00
0.77 0.00 0.11 0.70 0.06 0.04 0.22
down-regulated genes F1
0.00 0.00 0.00 0.00 0.00 0.00 0.00
0.60 0.00 0.48 0.26 0.00 0.00 0.00
0.75 0.00 0.79 0.75 0.00 0.00 0.01
0.93 0.00 0.93 0.61 0.00 0.00 0.01
0.79 0.00 0.66 0.90 0.00 0.00 0.01
0.79 0.00 0.53 0.51 0.00 0.00 0.01
0.77 0.00 0.67 0.76 0.00 0.00 0.02
0.81 0.00 0.55 0.67 0.00 0.00 0.02
0.95 0.00 0.57 0.89 0.00 0.01 0.04
0.77 0.00 0.57 0.86 0.00 0.00 0.04
0.71 0.00 0.38 0.47 0.00 0.00 0.07
0.74 0.00 0.82 0.70 0.00 0.00 0.01
0.60 0.00 0.72 0.59 0.00 0.00 0.01
0.74 0.00 0.80 0.50 0.00 0.00 0.04
0.95 0.00 0.49 0.61 0.00 0.00 0.03
0.64 0.00 0.41 0.58 0.00 0.00 0.06
0.63 0.00 0.50 0.36 0.00 0.01 0.02
0.45 0.00 0.61 0.57 0.00 0.00 0.02
0.73 0.00 0.80 0.87 0.00 0.00 0.06
0.63 0.00 0.60 0.48 0.00 0.00 0.17
0.84 0.00 0.77 0.70 0.00 0.01 0.01
knockout
reference
no-perturb
logFC-transfer
random-perturb
CAE
CPA
scPRAM

**Supplementary Figure 3. CrossSplit OOD evaluation results.** Heatmaps summarizing CrossSplit OOD evaluation results based on several common but less reliable metrics across the stimulated state of all gene knockouts of the Jurkat cell line in the Datlinger dataset. Each model was trained to predict the stimulated state of each cell type under the Out-of-Distribution (OOD) setting. Values represent the average performance across CrossSplit repetitions for baseline, deep learning and the reference models. Cases in which the reference model shows inferior performance compared to other models are highlighted in red.

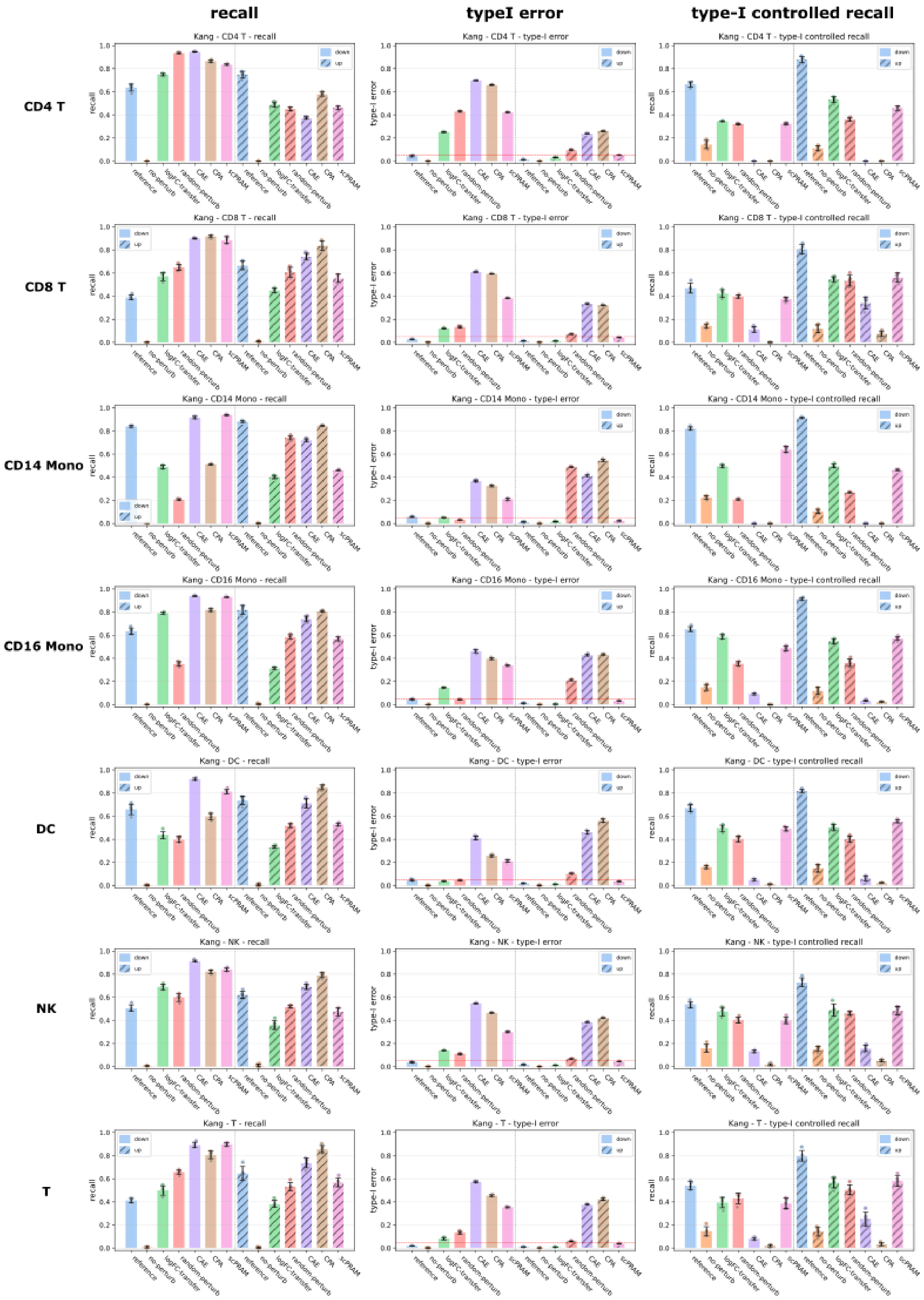
recall
typeI error
type-I controlled recall
CD4 T
CD8 T
CD14 Mono
CD16 Mono
DC
NK
T
Kang - CD4 T - recall
Kang - CD4 T - type-I error
Kang - CD4 T - type-I controlled recall
Kang - CD8 T - recall
Kang - CD8 T - type-I error
Kang - CD8 T - type-I controlled recall
Kang - CD14 Mono - recall
Kang - CD14 Mono - type-I error
Kang - CD14 Mono - type-I controlled recall
Kang - CD16 Mono - recall
Kang - CD16 Mono - type-I error
Kang - CD16 Mono - type-I controlled recall
Kang - DC - recall
Kang - DC - type-I error
Kang - DC - type-I controlled recall
Kang - NK - recall
Kang - NK - type-I error
Kang - NK - type-I controlled recall
Kang - T - recall
Kang - T - type-I error
Kang - T - type-I controlled recall
down
up
recall
type-I error
reference
no-perturb
logFC-transfer
random-perturb
CAE
CPA
scPRAM

**Supplementary Figure 4. Sensitivity analysis of models in identifying true DEGs for a variety of cell types in the Kang dataset**. Panels display recall values (left), type-I error (middle), and type-I-controlled recall (right), evaluated separately for up- and down-regulated DEGs. Recall correction is performed by adjusting the p-value threshold used for DEG detection, ensuring that the type-I error rate remains below 0.05. The recall of the reference model across most cell types is substantially below 1 and lower than that of most other models, especially for down-regulated DEGs. This behavior is driven by the high values of type-I error in most models, indicating that recall values in this setting are unreliable. Under controlled type-I error, the recall of the CPA model falls below that of most baselines, whereas scPRAM performs comparably to the baseline methods.

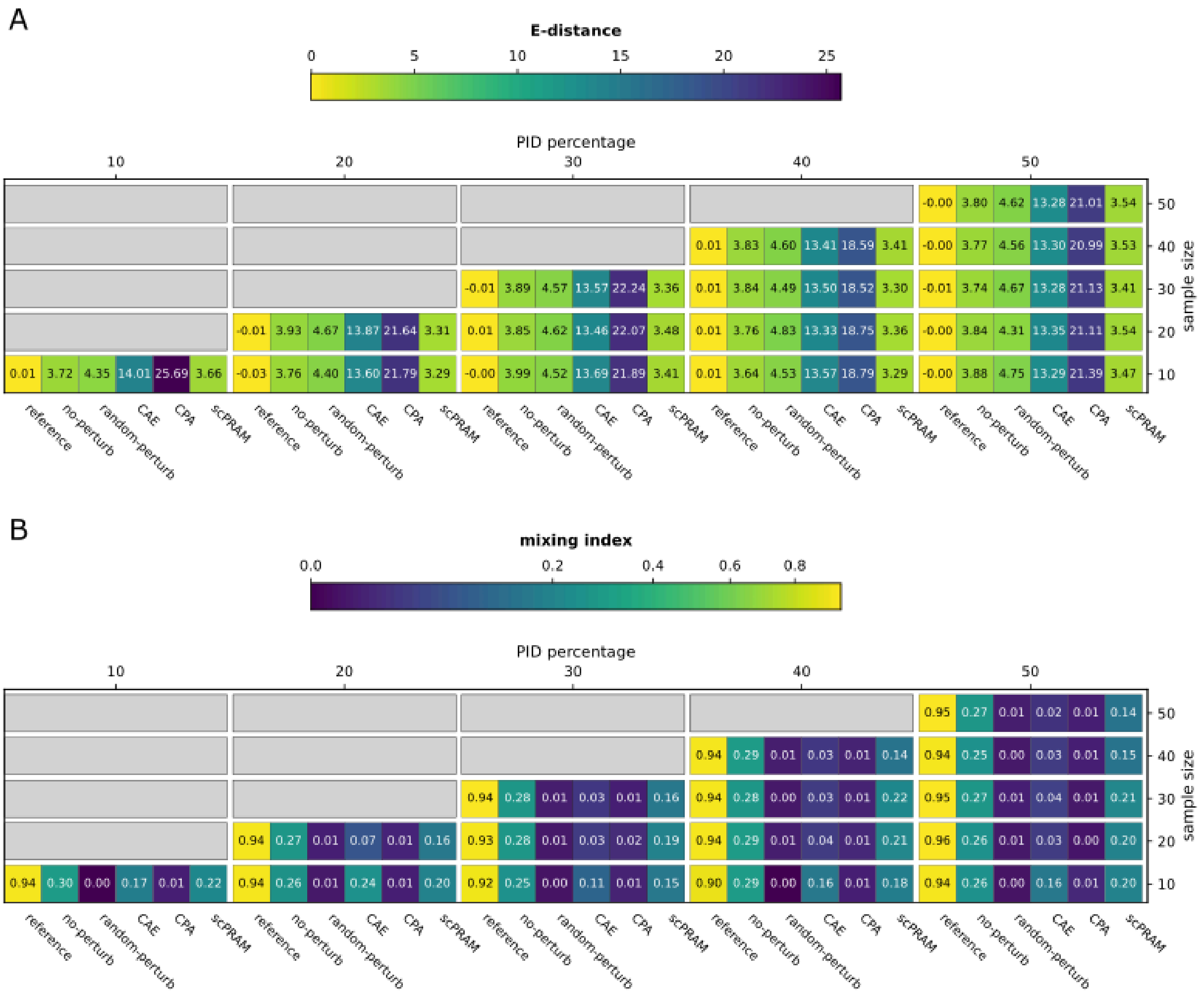


**Supplementary Figure 5. Model performance under the PID training setting across varying PID levels and samples sizes. (A)** E-distance and **(B)** mixing index for models trained with increasing proportions (10%, 20%, 30%, 40% and 50%) of perturbed B cells from the Kang dataset included during training. For each condition, performance is reported as the mean across five CrossSplit repetitions. At each PID (Partially In-Distribution) level, model predictions were compared with the evaluation (unseen) group using varying sample sizes (expressed as percentages), capped by the number of target perturbed cells that were '*seen'* during training, in order to avoid up-sampling the predictions of the reference (seen) model. For the mixing index heatmap, colors are scaled using a power-law normalization to enhance contrast among low-to-moderate values.

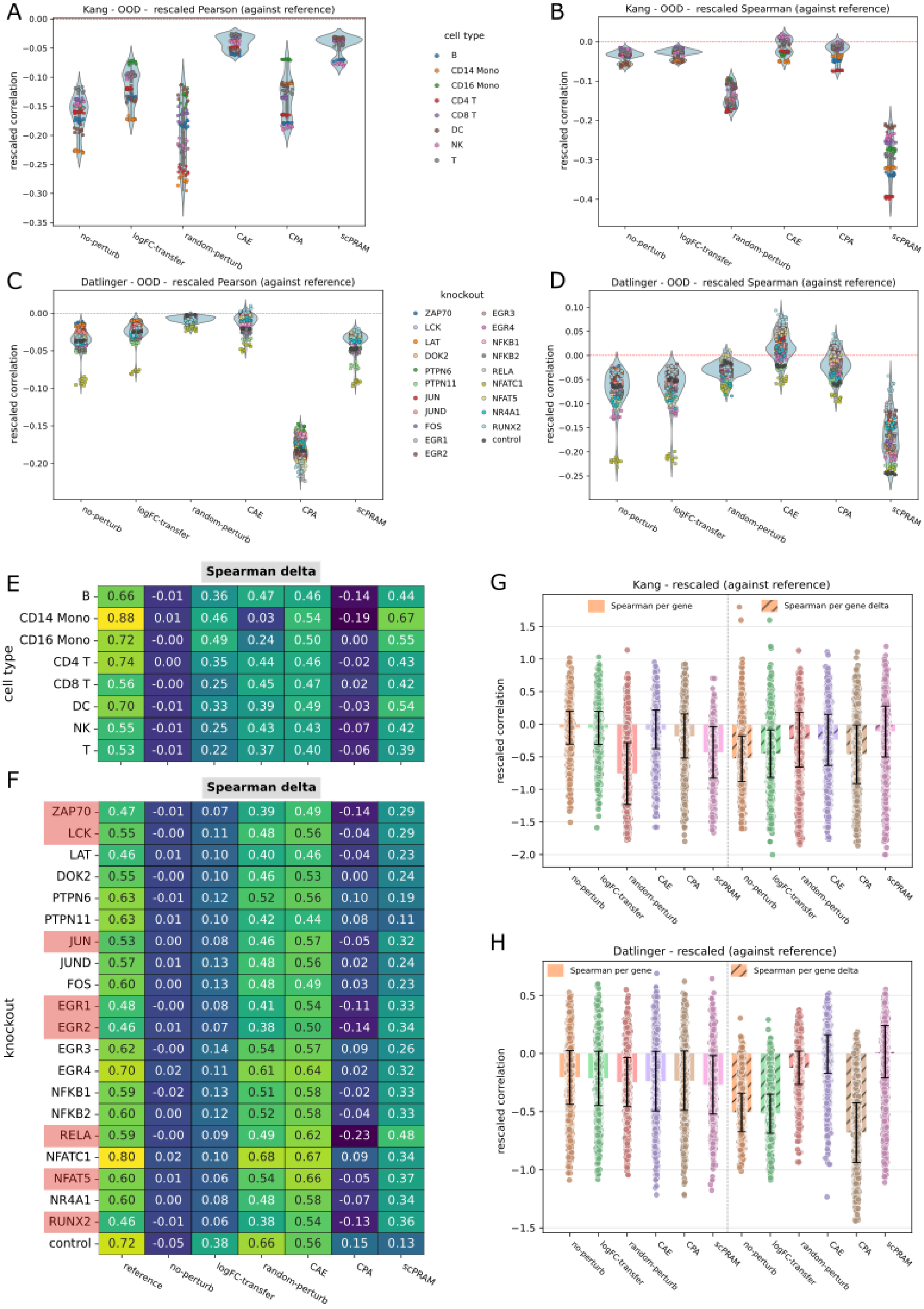
A
Kang - OOD - rescaled Pearson (against reference)
rescaled correlation
no-perturb
logFC-transfer
random-perturb
CAE
CPA
scPRAM
cell type
B
CD14 Mono
CD16 Mono
CD4 T
CD8 T
DC
NK
T
B
Kang - OOD - rescaled Spearman (against reference)
C
Datlinger - OOD - rescaled Pearson (against reference)
knockout
ZAP70
LCK
LAT
DOK2
PTPN6
PTPN11
JUN
JUND
FOS
EGR1
EGR2
EGR3
EGR4
NFKB1
NFKB2
RELA
NFATC1
NFAT5
NR4A1
RUNX2
control
D
Datlinger - OOD - rescaled Spearman (against reference)
E
Spearman delta
cell type
B 0.66 -0.01 0.36 0.47 0.46 -0.14 0.44
CD14 Mono 0.88 0.01 0.46 0.03 0.54 -0.19 0.67
CD16 Mono 0.72 -0.00 0.49 0.24 0.50 0.00 0.55
CD4 T 0.74 0.00 0.35 0.44 0.46 -0.02 0.43
CD8 T 0.56 -0.00 0.25 0.45 0.47 0.02 0.42
DC 0.70 -0.01 0.33 0.39 0.49 -0.03 0.54
NK 0.55 -0.01 0.25 0.43 0.43 -0.07 0.42
T 0.53 -0.01 0.22 0.37 0.40 -0.06 0.39
F
Spearman delta
knockout
ZAP70 0.47 -0.01 0.07 0.39 0.49 -0.14 0.29
LCK 0.55 -0.00 0.11 0.48 0.56 -0.04 0.29
LAT 0.46 0.01 0.10 0.40 0.46 -0.04 0.23
DOK2 0.55 -0.00 0.10 0.46 0.53 0.00 0.24
PTPN6 0.63 -0.01 0.12 0.52 0.56 0.10 0.19
PTPN11 0.63 0.01 0.10 0.42 0.44 0.08 0.11
JUN 0.53 0.00 0.08 0.46 0.57 -0.05 0.32
JUND 0.57 0.01 0.13 0.48 0.56 0.02 0.24
FOS 0.60 0.00 0.13 0.48 0.49 0.03 0.23
EGR1 0.48 -0.00 0.08 0.41 0.54 -0.11 0.33
EGR2 0.46 0.01 0.07 0.38 0.50 -0.14 0.34
EGR3 0.62 -0.00 0.14 0.54 0.57 0.09 0.26
EGR4 0.70 0.02 0.11 0.61 0.64 0.02 0.32
NFKB1 0.59 -0.02 0.13 0.51 0.58 -0.02 0.33
NFKB2 0.60 0.00 0.12 0.52 0.58 -0.04 0.33
RELA 0.59 -0.00 0.09 0.49 0.62 -0.23 0.48
NFATC1 0.80 0.02 0.10 0.68 0.67 0.09 0.34
NFAT5 0.60 0.01 0.06 0.54 0.66 -0.05 0.37
NR4A1 0.60 0.00 0.08 0.48 0.58 -0.07 0.34
RUNX2 0.46 -0.01 0.06 0.38 0.54 -0.13 0.36
control 0.72 -0.05 0.38 0.66 0.56 0.15 0.13
reference
no-perturb
logFC-transfer
random-perturb
CAE
CPA
scPRAM
G
Kang - rescaled (against reference)
Spearman per gene
Spearman per gene delta
rescaled correlation
H
Datlinger - rescaled (against reference)
Spearman per gene
Spearman per gene delta
rescaled correlation

**Supplementary Figure 6. CrossSplit OOD evaluation using correlation-based metrics. (A,B)** Violin plots with overlaid points showing CrossSplit evaluation results for the stimulated state of all cell types in the Kang dataset using **(A)** rescaled across-genes Pearson correlation and **(B)** rescaled across-genes Spearman correlation. Each point represents the result for a single random CrossSplit sampling for the specified cell type. Values are rescaled by subtracting the corresponding reference-model score. Therefore, values greater than zero indicate that the model outperforms the reference model. **(C,D)** Analogous results for the stimulated state of all gene knockouts of the Jurkat cell line in the Datlinger dataset, shown for **(C)** rescaled across-genes Pearson correlation and **(D)** rescaled across-genes Spearman correlation. **(E,F)** Delta variants of the across-genes Spearman correlation for the **(E)** Kang and **(F)** Datlinger datasets. Cases in which the reference model exhibits inferior performance relative to other models are highlighted in red. **(G,H)** Per-gene Spearman correlation (solid) and its delta variant (hatched), both rescaled relative to the reference model, shown for **(G)** Kang and **(H)** Datlinger datasets.

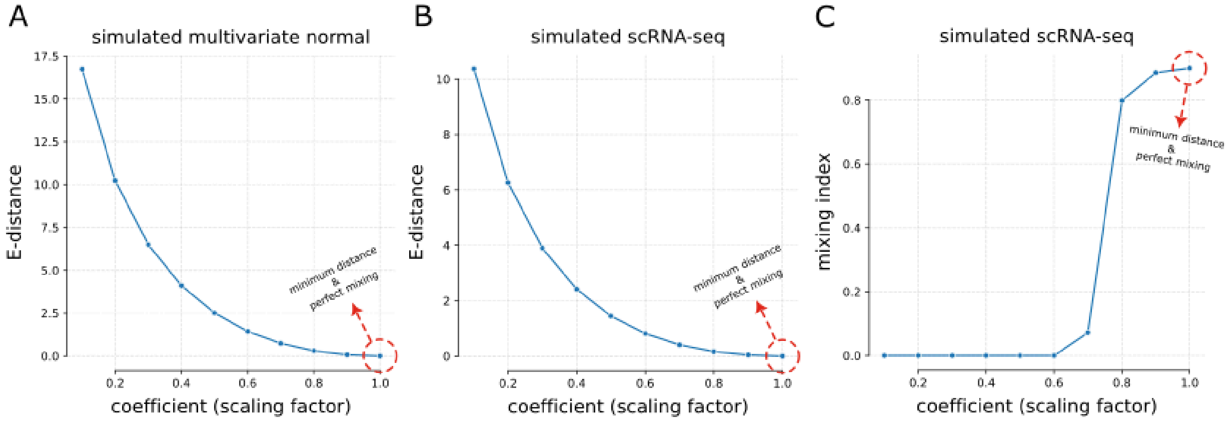


**Supplementary Figure 7. Evaluation of distribution-based metrics on simulated data.** Two samples with identical initial distributions were generated across 5,000 genes. Samples were drawn either from **(A)** a multivariate normal distribution with zero mean and identity covariance, or from **(B,C)** our single-seq RNA-seq simulator with read counts drawn from a negative binomial distribution (mean = 100, number of successes $r$ = 1). Initially, the samples were well mixed (coefficient = 1). The variance of one sample was then progressively reduced (coefficients 1 → 0.1). As expected, decreasing the variance of one sample increased the divergence between the two samples, as measured by **(A,B)** E-distance and **(C)** mixing index.

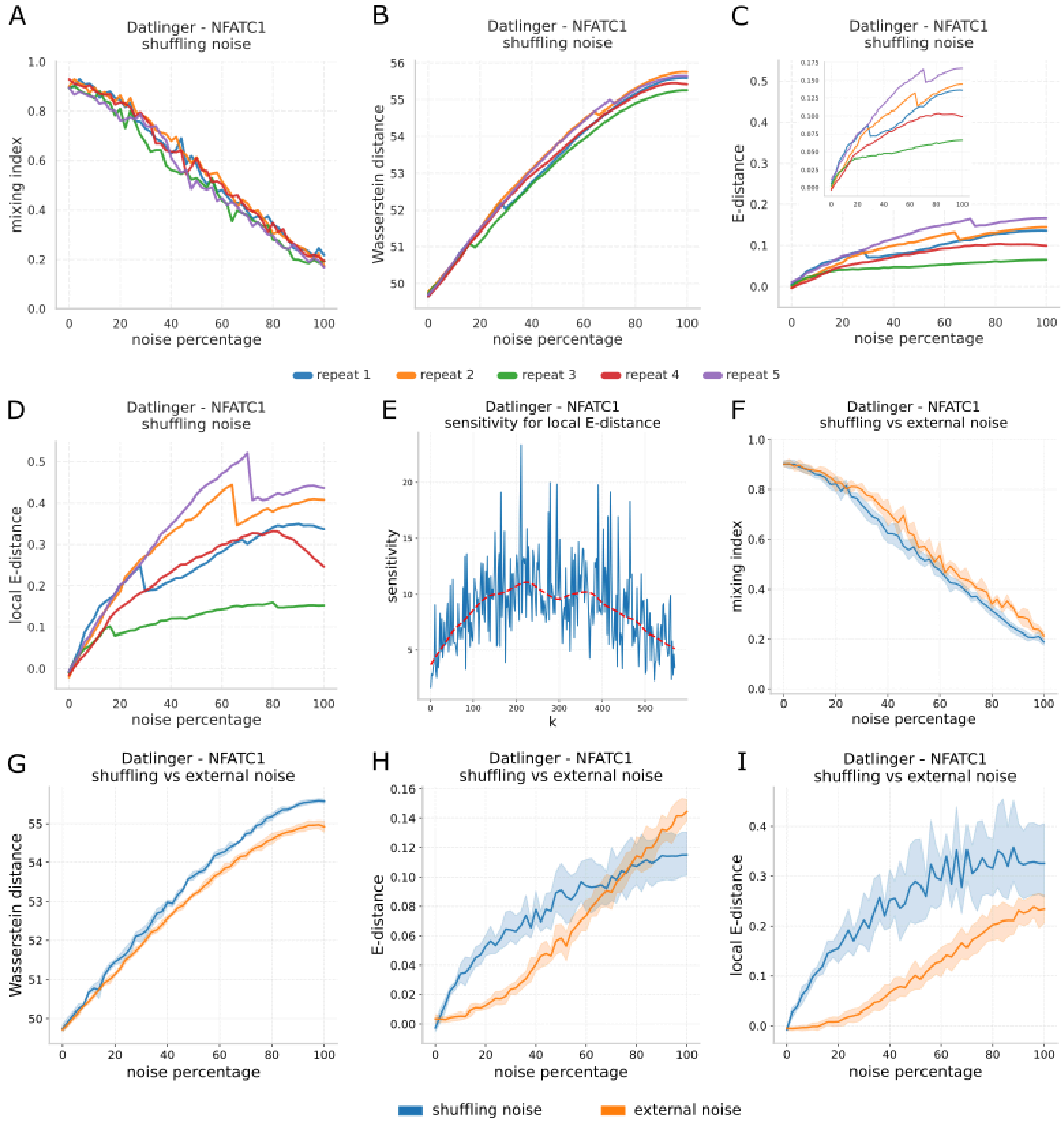


**Supplementary Figure 8. Performance of distribution-based metrics in capturing gene-gene interactions and matrix-level noise. (A-D)** Values of **(A)** mixing index, **(B)** Wasserstein distance, **(C)** E-distance and **(D)** local E-distance under nested shuffling-noise experiment, computed between the noisy and test sets of stimulated NFATC1-knockout Jurkat cells from the Datlinger dataset across noise levels ranging from 0% to 100%. **(E)** Sensitivity of local E-distance to shuffling-induced disruptions across different neighborhood sizes ($k$) in NFATC1-knockout Jurkat cells, with the maximum value observed at $k \approx 225$. **(F-I)** Comparing the effect of non-nested shuffling-noise and matrix-level noise-addition (external noise) on stimulated NFATC1-knockout Jurkat cells, measured by **(F)** mixing index, **(G)** Wasserstein distance, **(H)** E-distance and **(I)** local E-distance. Shaded regions denote variability across five repetitions.

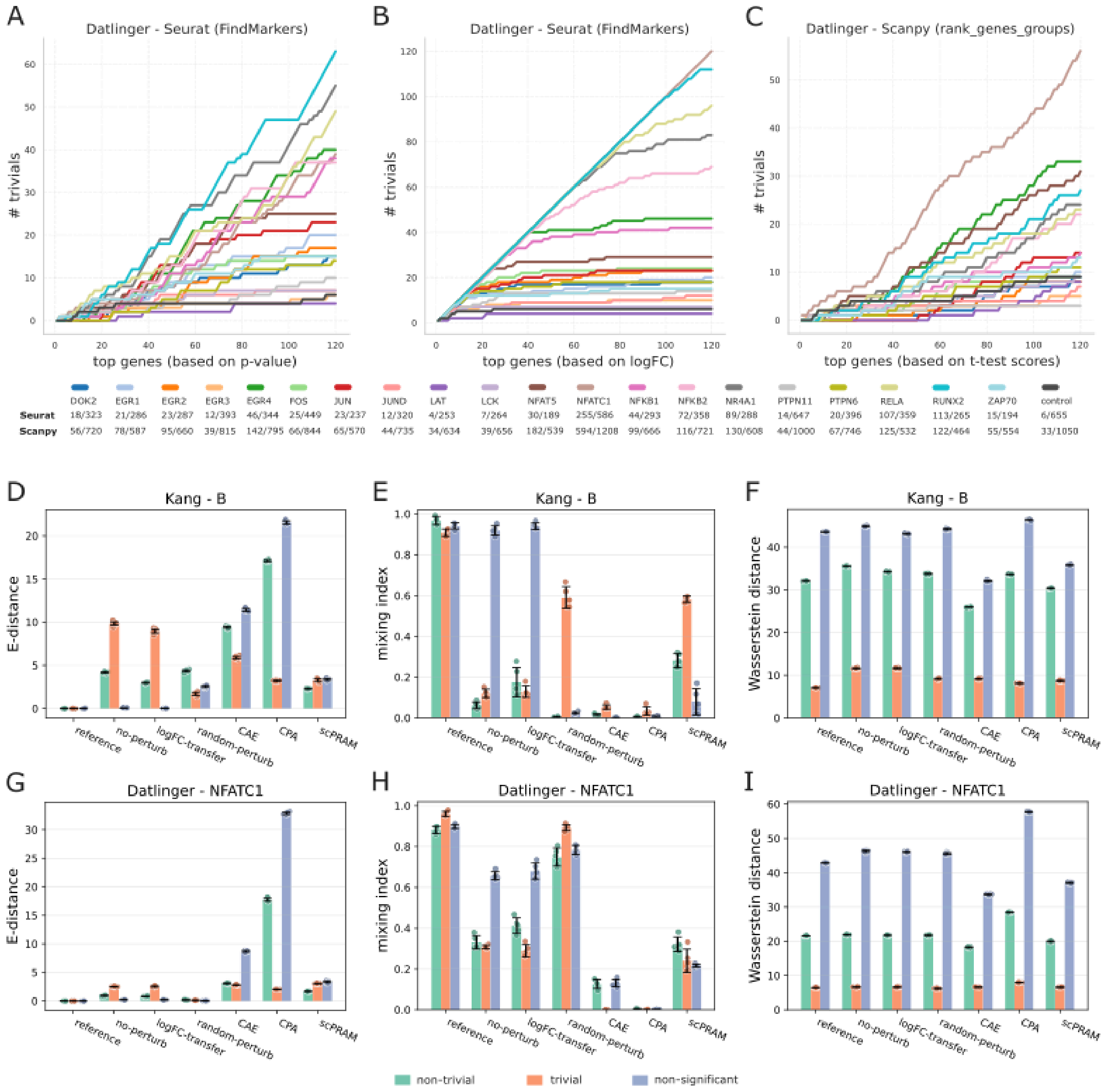


**Supplementary Figure 9. CrossSplit evaluation of model performance across gene categories. (A-C)** Prevalence of trivial genes among top-ranked DEGs in the Datlinger dataset, ranked by **(A)** p-value, **(B)** logFC among significant genes obtained using Seurat's *FindMarkers* function, and **(C)** absolute z-scores computed using Scanpy's *rank_genes_groups* function with a t-test. **(D-F)** CrossSplit evaluation of models on trivial, non-trivial and non-significant genes in stimulated B cells from the Kang dataset using: **(D)** E-distance, **(E)** mixing index, and **(F)** Wasserstein distance. **(G-I)** Corresponding CrossSplit evaluations for stimulated NFATC1-knockout Jurkat cells from the Datlinger dataset, using the same gene categories and metrics: **(G)** E-distance, **(H)** mixing index, and **(I)** Wasserstein distance.

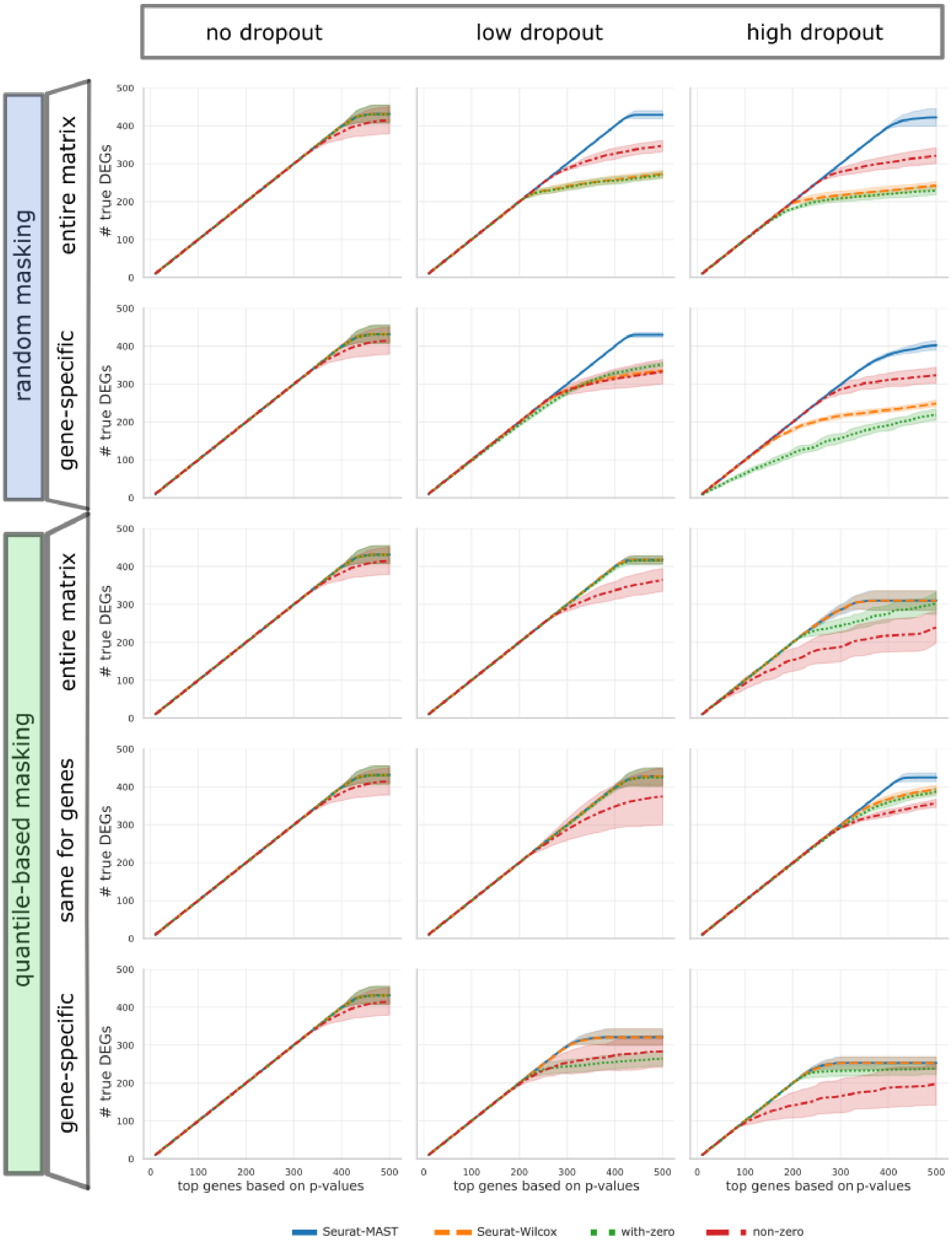

no dropout
low dropout
high dropout
random masking
entire matrix
gene-specific
quantile-based masking
entire matrix
same for genes
gene-specific
### true DEGs
top genes based on p-values
Seurat-MAST
Seurat-Wilcox
with-zero
non-zero

**Supplementary Figure 10. Comparison of the power of differential expression detection under varying dropout scenarios.** Shown is the number of true differentially expressed genes (DEGs) recovered among the top-ranked genes identified by different statistical methods on simulated single-cell perturbation data. Each approach computed p-values for differential expression between perturbed and unperturbed cells. For each method, genes were ranked by statistical significance (perturbed vs. unperturbed), and recovery of true DEGs was assessed among the top-ranked genes. Results are shown under three dropout probabilities: no dropout, low dropout ($P_{dropout} = 0.8$), and high dropout ($P_{dropout} = 0.93$), resulting in final zero-expression proportions of 0.37, 0.87 and 0.95, respectively. The maximum dropout probability was selected such that the resulting sparsity closely matched that of the real Kang dataset (approximately 95% zero values). For low and high dropout, five masking mechanisms were evaluated, including two random-based strategies (all-genes, gene-specific) and three quantile-based strategies (all-genes, same, gene-specific). The no-dropout (no-mask) condition is identical across masking scenarios and is therefore repeated across rows for visual comparison. As shown, increasing dropout probability generally reduced DEG-recovery performance, although the extent of the decline varied across methods and masking schemes. Notably, Seurat-MAST consistently outperformed the other approaches across dropout scenarios, demonstrating superior robustness to high sparsity.

#### References


1. Jiang, R., Sun, T., Song, D. & Li, J. J. Statistics or biology: the zero-inflation controversy about scRNA-seq data. *Genome Biol* **23**, 31 (2022).
2. Finak, G. *et al.* MAST: a flexible statistical framework for assessing transcriptional changes and characterizing heterogeneity in single-cell RNA sequencing data. *Genome Biol.* **16**, 278 (2015).
